# Fatigue-related changes in value-based decision-making during listening are associated with increased frontal cortical activation

**DOI:** 10.64898/2026.09.11.750884

**Authors:** Alina Schulte, Cora Jirschik Caron, Dorothea Wendt, Ingrid Johnsrude, Katarzyna Dudzikowska, Selma Lugtmeijer, Matthew Apps, Jörg Encke, Torsten Dau, Hamish Innes-Brown

**Affiliations:** Eriksholm Research Center, Oticon A/S, Snekkersten, Denmark; Hearing Systems Section, Department of Health Technology, Technical University of Denmark, Kongens Lyngby Denmark; Centre for Human Brain Health, School of Psychology, University of Birmingham, United Kingdom; Department of Psychology & Brain and Mind Institute, The University of Western Ontario, London, Ontario, Canada; Macquarie University, Department for Linguistics, Sydney, Australia

**Keywords:** Listening-related fatigue, decision-making, effort-discounting, functional near-infrared spectroscopy, hearing loss

## Abstract

Hearing aid users commonly report listening-related fatigue, which may contribute to disengagement from demanding listening situations and negative psychosocial outcomes. Hearing rehabilitation currently lacks strategies to mitigate listening fatigue, partly because fatigue build-up and its consequences are difficult to measure beyond self-report.

This study investigated whether value-based listening decisions and associated frontal cortical activity are modulated by fatigue. Thirty hearing aid users completed a one-hour listening paradigm while cortical hemodynamic activity was measured using functional near-infrared spectroscopy (fNIRS). On each trial, participants decided whether to engage in listening or rest based on an offer defined by expected task difficulty and reward, and rated their momentary fatigue.

Fatigue ratings increased over time, and unsuccessful listening trials were associated with larger subsequent increases in fatigue, suggesting that performance contributed to changes in self-reported fatigue. Listen-versus-rest decisions followed the expected effort-reward trade-off, with participants more likely to accept offers that provided higher rewards and lower listening demands. Over the course of the experiment, participants became less likely to accept listening offers; however, neither time on task nor subjective fatigue altered the weighting of reward and difficulty. In contrast, frontal cortical activation during evaluation of accepted offers increased over time for high-difficulty offers with low or medium rewards. Follow-up analyses indicated that frontal oxygenation and listening decisions were more strongly associated with time on task than with subjective fatigue ratings.

Together, these findings suggest that behavioral and neural measures obtained before listening capture fatigue-related processes beyond self-report, providing a basis for identifying and potentially addressing fatigue in hearing rehabilitation.

## Introduction

Individuals with hearing loss are thought to compensate for degraded acoustic input by allocating additional cognitive resources to understand speech, particularly in noisy environments (Downs, 1982; Pichora-Fuller et al., 2016; Rakerd et al., 1996). Hearing loss not only makes speech-in-noise understanding more effortful, but sustained listening effort may also increase the need for recovery and reduce overall well-being (Holman et al., 2021, Naachtegal et al., 2009; Shinn-Cunningham et al., 2008; Rönnberg et al., 2013)).

Listening-related fatigue can be understood as a subjective feeling of tiredness, exhaustion, or lack of energy that arises when sustained listening demands exceed available cognitive or motivational resources (Bess and Hornsby, 2014; Schneider et al., 2019). Within the Framework for Understanding Effortful Listening (FUEL, Pichora-Fuller et al., 2016), effort investment is conceptualized as a function of both task demands and motivation. This perspective also aligns with broader theoretical accounts of effort, motivation, and cognition, which propose that individuals allocate limited cognitive resources according to task difficulty and goal importance (Brehm and Self, 1989; Hockey, 2013; Kahneman and Tversky, 1973; Richter et al., 2016). Across these accounts, sustained effort is maintained as long as it is perceived as worthwhile and effective. Fatigue is thought to arise when sustained effort does not sufficiently contribute to achieving the intended goal, reflecting an unfavorable effort-reward balance (e.g., unsuccessful communication despite high cognitive effort, see also Figure 2 in Davis et al., 2021).

Generally, fatigue serves as an indicator of the need for breaks and plays an important role in enabling recovery, thereby helping to regulate psychological resources and support continued goal-directed behavior (Hockey, 2011). Among individuals with hearing loss, however, fatigue is often experienced not as an episodic state but as a chronic burden that can manifest in physical, mental, emotional, or social forms (Davis et al., 2021). When sustained over longer periods, such ongoing fatigue can adversely affect psychosocial functioning and perceived hearing handicap (Alhanbali et al., 2018; Hornsby and Kipp, 2016), potentially contributing to broader negative health outcomes and reduced well-being.

Evidence regarding the influence of hearing aids on listening-related fatigue is mixed (Davis et al., 2021; Holman et al., 2020), suggesting that their effects are complex and may vary across contexts and individuals. For example, Blümer et al. (2024) who assessed listening-related fatigue over longer time periods using a “time-compressed acoustic day” –paradigm, and Holman et al. (2021), who employed a longitudinal design before and after hearing aid fitting, both reported reductions in fatigue related to hearing aid use. In contrast, in a study in which hearing aid users performed a dual-task paradigm in both aided and un-aided conditions, the aided condition resulted in better word recall and shorter reaction times but did not reduce subjective fatigue (Hornsby, 2013). Consequently, reductions in fatigue are not consistently observed following hearing aid intervention, leaving listening-related fatigue a common and largely unaddressed complaint. In some cases, removing the hearing aids has been reported as a strategy to disengage from listening and alleviate acute overstimulation (Cumming et al., 2026). No well-established clinical approaches currently exist to address listening fatigue directly. For hearing healthcare to better support the management of listening-related fatigue, a deeper understanding of how fatigue develops over time, physiological correlates, and the cognitive and motivational processes that precede its build-up is required.

Investigating fatigue build-up is not trivial, as conscious awareness of fatigue may lag behind its underlying physiological and cognitive processes, making fatigue difficult to recognize and manage in everyday life and complicating its assessment using subjective measures alone. To capture listening-related effort and fatigue, a range of measures have been proposed, including self-report questionnaires (Alhanbali et al., 2017; Krueger et al., 2017), behavioral indices such as reaction times (Houben et al., 2013), dual tasks (Gosselin and Gagné, 2010), and physiological markers such as skin conductance, heart-rate variability (Mackersie and Calderon-Moultrie, 2016) and pupil dilation (Ohlenforst et al., 2017; Wang et al., 2018; Zekveld et al., 2010). Neural metrics that have been associated with fatigue are electroencephalographic alpha power (e.g. Dimitrijevic et al., 2019) and frontal cortex hemodynamics (e.g. Rovetti et al., 2022; see also Peelle, 2018). However, these outcome measures are often only weakly correlated with each other, demonstrating the complexity of these constructs, and suggesting that they either capture different aspects of listening-related effort and fatigue or reflect differences in how these constructs are operationalized and measured (Alhanbali et al., 2019; McGarrigle et al., 2014).

Although motivational factors have been identified as dynamic contributors to listening effort and listening-related fatigue, most studies have treated motivation as constant and have not manipulated them explicitly in listening experiments. Only recently, studies have utilized decision-making tasks in which participants can choose whether to make an effort in exchange for reward, or to take a rest (Matthews et al., 2023; McLaughlin et al., 2021). This type of task allows direct examination of how choices to work or rest depend on task difficulty and reward, and how this trade-off changes as fatigue accumulates. Using such a task, effort-reward evaluation was found to result in a reduced willingness to exert effort as fatigue accumulates (Matthews et al., 2023; Müller et al., 2021). Interestingly and in line with Hockey (2013), making mistakes, such that the participant failed to achieve the intended goal, was associated with additional increases in reported fatigue (Matthews et al., 2023). While fatigue has been shown to influence effort-reward evaluations in physical and arithmetic-task paradigms (Matthews et al., 2023; Müller and Apps, 2019), it is currently unclear whether listening-related fatigue similarly influences effort-reward evaluations during speech perception.

Previous work has employed computational models that decompose the fatigue into recoverable, short-term and unrecoverable, longer-term components and provide a trial-by-trial estimate of the subjective value of working (Matthews et al., 2023; Müller et al., 2021). Müller et al. (2021) used fMRI to identify brain areas in which BOLD responses covaried with these model-derived fatigue components during effort-reward evaluation. Specifically, the cingulate cortex and middle frontal gyri were associated with longer-term fatigue build-up, a posterior region in the cingulate cortex with shorter-term fluctuations (recoverable fatigue), and BOLD responses in the superior frontal gyri and ventral striatum with the current subjective value of working.

Building on previous research, we adapted existing effort-discounting paradigms that manipulate both reward and task difficulty (Matthews et al., 2023; Müller et al., 2021) to a speech-in-noise intelligibility task. Presented as a listening game with the goal of earning as many points as possible, participants chose on each trial whether to engage in a listening task, in which they repeated a sentence presented in noise, or to take a short rest. Each choice was preceded by an offer specifying both the listening difficulty and the number of points that could be earned if the sentence was repeated correctly. This design allowed us to examine whether accumulating fatigue changes how hearing-aid users evaluate the trade-off between listening effort and expected reward. In addition to decision behavior, we employed functional near-infrared spectroscopy (fNIRS) to investigate changes in cortical oxygenation during offer evaluation as fatigue developed. Regions-of-interest focused on middle and superior frontal gyri given their association with unrecoverable fatigue and subjective value in Müller et al. (2021).

Both decision making and mental fatigue have repeatedly been related to changes in oxygenation in frontal brain regions, which can be measured using functional near-infrared spectroscopy (fNIRS) (Hamann & Carstengerdes, 2023; Lin et al., 2019; Nihashi et al., 2019; Skau et al., 2019; Varandas et al., 2022; see Yan et al. (2025) for a review). For example, during cognitive tasks such as the Stroop task, subjective fatigue has been associated with declining prefrontal cortex activity over time (Nihashi et al., 2019), as well as with reduced activity in individuals with traumatic brain injury compared to healthy controls (Skau et al., 2022). This may reflect reduced cognitive control and sustained attention during mental fatigue (Ishii et al., 2014). fNIRS studies on decision-making have used e.g. the Balloon Analogue Risk Task (Cazzell et al., 2012), Iowa Gambling task (Li et al., 2019) or Ultimatum Game (Vanutelli et al., 2020); these studies have consistently implicated prefrontal cortical activity in decision evaluation and risk assessment.

fNIRS is well suited for auditory tasks in participants with hearing devices, as it is quiet and is not affected by hearing-device-related artefacts in the same way as fMRI. Although previous fNIRS studies have linked higher prefrontal cortex oxygenation to higher listening demands (Rovetti et al., 2021, 2022; Vaisberg et al., 2024; see Shatzer & Russo (2023) for a review), the impact of listening-related fatigue on cortical activation during value-based listening decisions remains unexplored.

We therefore examined whether listening-related fatigue predicts trial-by-trial decisions to engage in a listening task or take a rest under different combinations of task difficulty and reward, and whether these decision-related effects are reflected in frontal cortical oxygenation.

### We tested three broad hypotheses

First, we expected fatigue ratings to increase over the course of the task, and trial-to-trial changes in fatigue to be modulated by difficulty level and exerted effort, reward incentives, the listening-versus-rest decision, and trial success.

Second, we hypothesized that the choice to listen versus rest would reflect the difficulty-reward trade-off specified by each offer. Additionally, we expected this evaluation to change over time, such that participants would become less likely to accept offers involving high difficulty and low reward. This would suggest a change in the subjective valuation of effort relative to reward as fatigue accumulated.

Third, we expected frontal cortical oxygenation during offer evaluation to change as fatigue accumulated, particularly for offers requiring greater effort relative to reward, indicating altered neural processing of the decision to engage in listening before auditory stimulation occurred.

## Methods

### Participants

Thirty native Danish-speaking hearing aid users with sensorineural hearing loss were recruited from the test participant database at Eriksholm Research Center (14 females, 16 males; mean age = 73.7 years, SD = 7.1 years). All participants were regular users of Oticon More or Oticon Real (Oticon A/S, Smørum, Denmark) hearing aids. Four-frequency pure-tone average hearing thresholds across both ears (PTA4; 0.5, 1, 2, 4 kHz) ranged from 28.1 to 73.8 dB HL (mean = 47.8, SD = 12.8), with a maximum difference in PTA4 between ears of 11.25 dB HL (mean = 4.0 dB HL, SD = 2.8 dB HL).

Participants were drawn from a larger sample enrolled in a previous field trial on listening-related fatigue (Micula et al., 2026). Selection aimed to include individuals who showed within-person variation in fatigue ratings during the field trial, suggesting that they experienced fluctuations in listening-related fatigue in everyday life and were able to report these changes reliably. Additional considerations included hearing-aid model compatibility and minimizing age-related variability.

**Figure 1.**
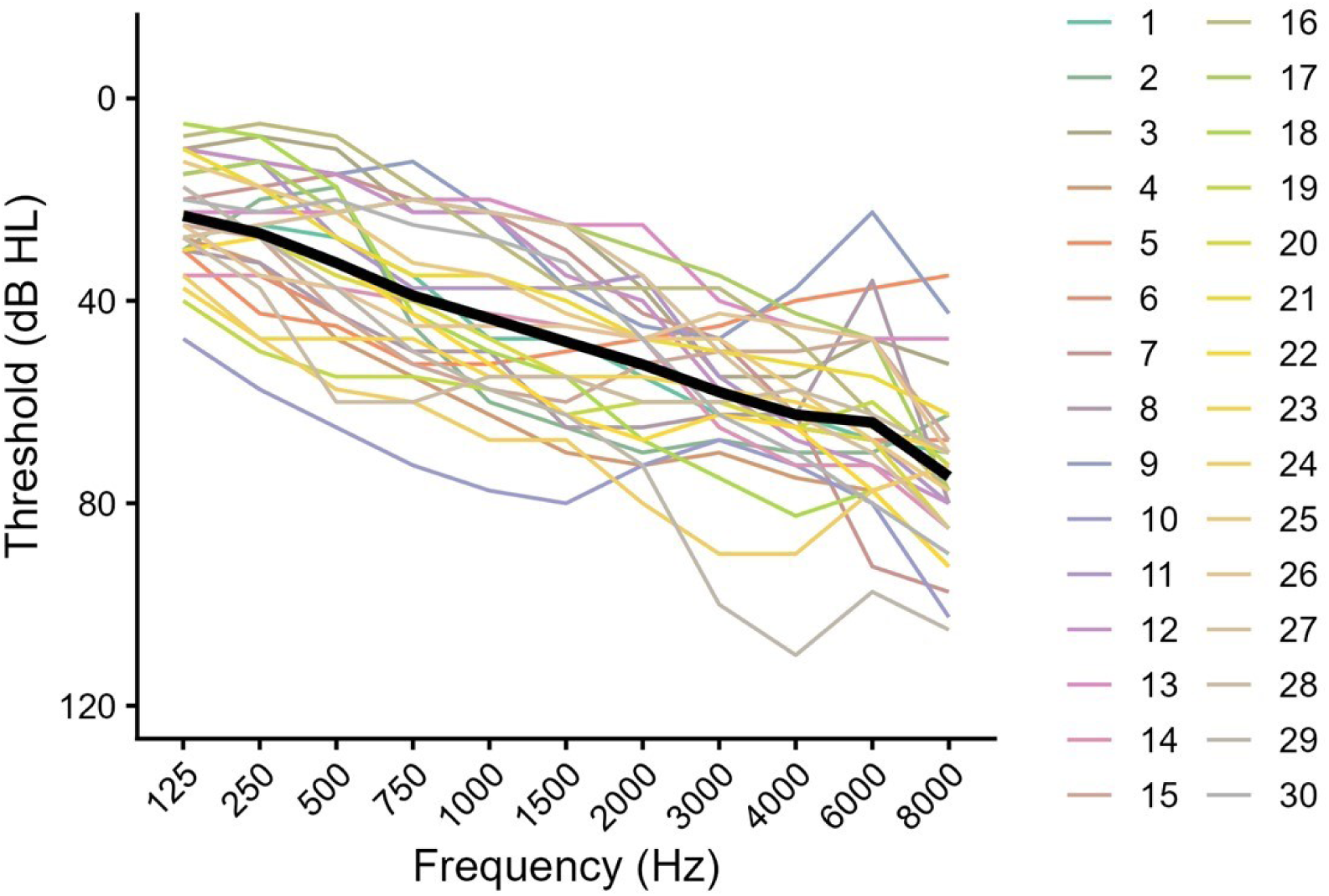
Audiograms. Pure-tone audiometry thresholds were obtained at 0.125, 0.25, 0.5, 0.75,1, 1.5, 2, 3, 4, c and 8 kHz and averaged across both ears.

Participants were fitted with test hearing aids for the visit based on the most recent audiogram. Individual haring-aid settings were kept as similar as possible to those of the participants’ own devices, except that all additional noise reduction features were turned off (i.e. no beamforming or sudden sound stabilizer).

All participants gave written consent. Ethical approval was obtained from the Danish Science-Ethics Committee (reference H-16036391).

### Lab setup

The study was conducted in a sound-treated booth at Eriksholm Research Center, Denmark. Speech-in-noise stimuli were presented from a frontally positioned loudspeaker at head height (Genelec 8030C). Auditory stimuli were presented using custom MATLAB scripts for the SRT estimation task and PsychoPy in Python for the effort-discounting paradigm. Stimuli were played via a computer and routed through an RME Madiface soundcard with a Ferrofish Pulse 16 I/O extension. Visual stimuli were shown on a standard desktop monitor beneath the loudspeaker. Additionally, eye tracking data and a video recording of the participants’ faces were collected and analyzed as part of a separate study^1^.

Cortical oxygenation changes were recorded using an fNIRS system (NIRx NirScout) with 16 LED sources and 16 avalanche photodiode detectors. Near-infrared light at 760 nm and 850 nm wavelengths was sampled at a rate of 3,9063 Hz using the NIRStar 15.3 acquisition software (NIRx Medizintechnik GmbH, Berlin, Germany).

### fNIRS montage

The fNIRS montage targeted superior and middle frontal gyri, regions implicated in mental fatigue and value-based decision-making (Vanutelli et al., 2020). Activity in these regions has also been associated with computationally derived estimates of unrecoverable fatigue and subjective value in a comparable effort-discounting paradigm (Müller et al., 2021). Using the fOLD toolbox (Zimeo Morais et al., 2018), channels with at least 38% anatomical specificity to bilateral superior and middle frontal gyri were identified and combined into a single frontal ROI.

**Figure 2.**
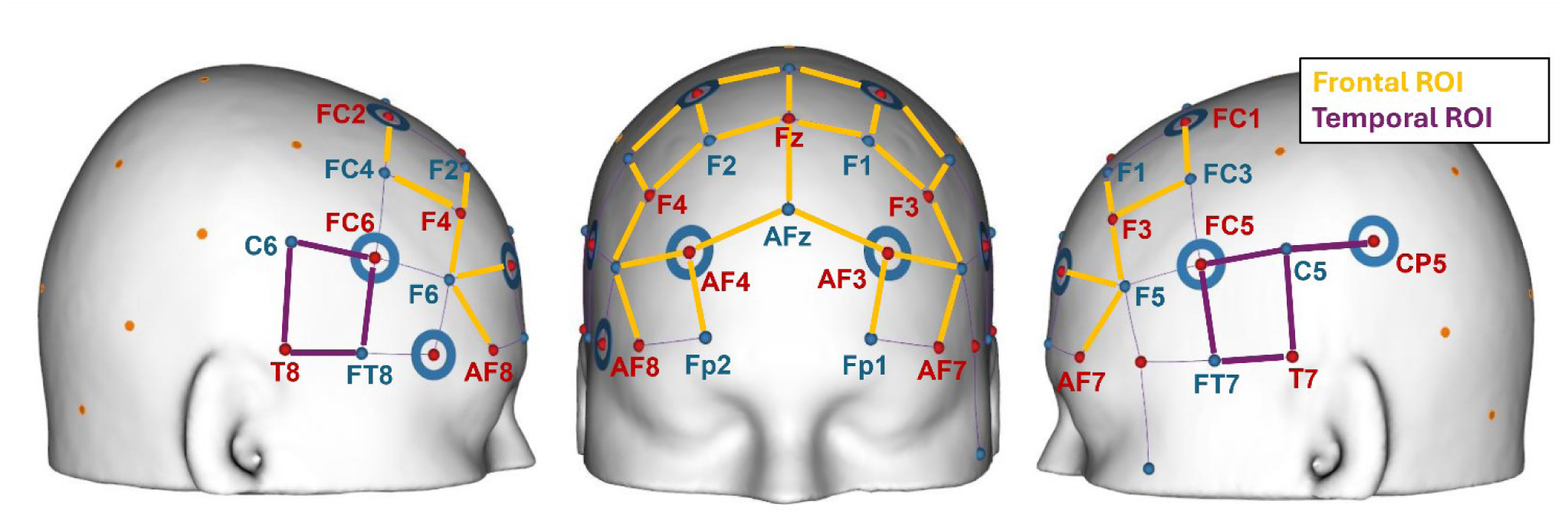
fNIRS montage. The frontal region-of-interest (ROI) targeting bilateral middle-and superior frontal gyri included the following source-detector pairs (in 20-20 system notation): Fp1-AF3, AF3-F5, AF7-F5, F3-F5, F3-F1, F3-FC3, FC1-F1, Fz-AFz,Fz-FCz, FCz-FC2, FC2-F2, FC2-FC4, F4-F2, F4-FC2, F4-Fc, AF4-AFz, AF4-Fp2, AF4-Fc and AF8-Fc, as indicated in yellow. Channels indicated in purple (FC5-FT7, FC5-C5, T7-C5, T7-FT7, FCc-Cc, FCc-FT8, T8-FT8, T8-Cc) refer to temporal ROI (control region), targeting bilateral superior temporal gyri.

Additionally, the montage targeted an auditory cortical region that served as a control ROI, as it was not expected to be directly involved in decision-making or fatigue-related processes. This auditory region ROI predominantly targeted the superior temporal gyri, with additional coverage of parts of the middle temporal gyri and inferior frontal gyri. Corresponding channel pairs were selected in NIRSite using standard 10-20 optode locations (see **Figure 2**) and implemented using an NIRx cap matched to each participant’s head size and aligned to standard anatomical landmarks (nasion, inion, and preauricular points).

### Listening-effort discounting task

A traditional speech intelligibility test, in which participants are presented with speech-in-noise and instructed to repeat the heard sentence as accurately as possible, was transformed into a listening game, where the participant’s task was to earn as many points as possible during the entire experiment. In each trial, participants were presented with an offer showing the difficulty level (easy, medium or hard) together with the number of points the participant could earn (either 6,8 or 10) if they chose to *listen* and successfully repeated the coming sentence correctly. Reward levels and difficulty levels were fully crossed, such that each reward value was paired with each difficulty level, yielding nine unique offer conditions. If participants chose to *rest*, they were shown a visually neutral image depicting a landscape or city scene and no additional sound. Rest would always give 1 point (**Figure 3**).

**Figure 3.**
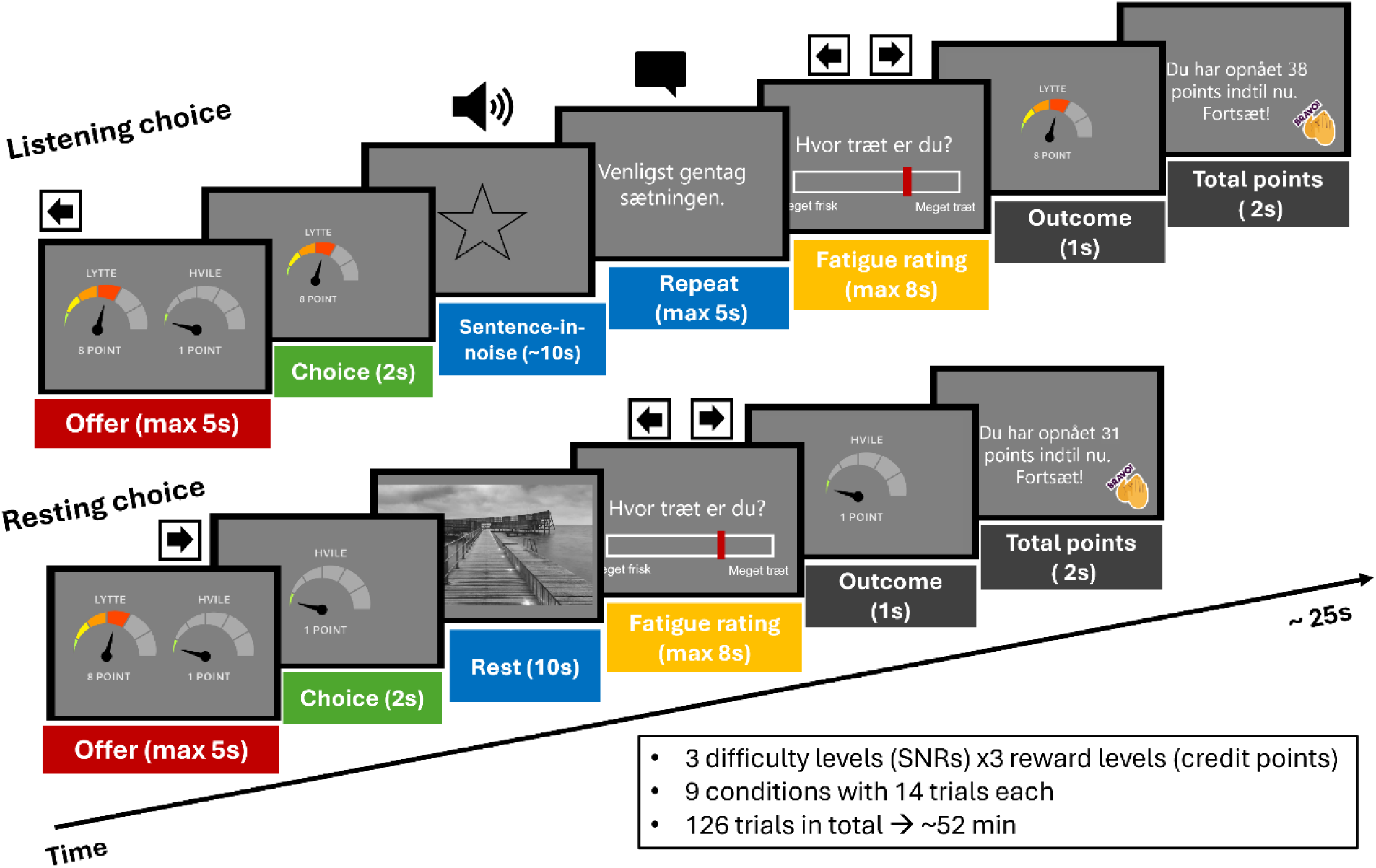
Trial procedure. Schematic representations of a listening trial (top) and a rest trial (bottom). Participants are presented with an offer consisting of a reward level and listening difficulty and opt either to listen or to rest. On listening trials, participants listened to a sentence presented in noise, verbally repeated the sentence, rated their current fatigue, received feedback on the trial success and their cumulative score. On rest trials, the sentence-in-noise and repetition phases were replaced by a 10-s rest period during which participants viewed a scenery image without auditory stimulation. There was no feedback, and the cumulative score was the previously presented score, plus one. Arrows indicate the response keys used for option selection and fatigue ratings. Display durations for the offer and fatigue rating screens represent maximum durations; both screens could be advanced by a button press

If no decision was made within 5 s, the text “Missed trial. Please try to choose faster.” was displayed in Danish before the next offer was presented.

On each non-missed trial, following either the listening or *rest* period and, for listening trials, the verbal repetition of the sentence, participants were asked to rate how mentally tired they currently felt. A scale from 0 to 100 appeared on the screen, with the cursor remaining in the previously chosen position. Participants had up to 8 s to indicate their momentary subjective fatigue in response to the question “How tired are you?” presented in Danish. Participants were instructed to interpret tiredness as mental or cognitive fatigue specifically arising from listening and concentrating on the task, rather than general sleepiness. Arrow keys were used to move the cursor from the previous trial’s rating in steps of 5 (left or right) or 1 (up or down). A baseline fatigue rating was obtained at the beginning of the experiment without a time limit.

### Scoring Rule and Feedback

Each sentence contained three keywords. Verbal responses were scored online by the experimenter as correct if at least two of the three keywords were repeated correctly or incorrect otherwise, which determined the feedback presented to the participant. If a response was correct, the cumulative score was the previous score plus the reward for that trial. If fewer than two keywords were repeated correctly, the participant did not receive any reward points and the cumulative score remained the same as for the previous trial. Minor variations in participants’ responses (e.g., dialectal variations, self-corrections, or the addition of extra words) were accepted as long as the sentence meaning was preserved, whereas grammatical changes, such as changes in verb tense, were scored as incorrect (Kressner et al., 2026).

### Stimuli

#### Listening stimuli: Speech-in-noise sentences

Sentence stimuli were taken from the Danish Auditory Sentence test (DAST; Kressner, 2023), using the female speaker F1. From the 1200 available sentences in the auditory DAST material we selected 126 sentences (9 conditions x 14 trials = 126 sentences) that had received the highest naturalness ratings (Kressner et al., 2024) and whose psychometric-function inflection points were closest to the corpus average (Kressner et al., 2025), thereby minimizing differences in intelligibility between sentences.

Background noise consisted of 16-talker babble created by overlaying 16 independently shuffled versions of the concatenated F1 sentences. Using the same sentence corpus for both the babble and target speech ensured spectral similarity between masker and target. In total, 42 independent noise segments were extracted from the continuous 16-talker babble. For each trial, a randomly selected babble noise segment was trimmed to match the duration of that trial’s target sentence, including 3 s preceding and following the sentence, gated with a 1 s linear onset and offset ramps, and RMS-normalized to an RMS level of −26 dB relative to full scale.

Noise levels were kept constant and sentence level was adjusted to achieve a target signal-to-noise ratio (SNR) based on each participant’s estimated speech reception threshold at 50% intelligibility (SRT50; see Procedure). The amplitude of each sentence was further corrected using sentence-specific offset values provided with the stimulus material to account for intelligibility differences between sentences.

For each participant, 42 sentences were prepared for each difficulty level. Easy, medium, and hard corresponded to target SNRs of SRT₅₀ + 8 dB, SRT₅₀ + 3 dB, and SRT₅₀ − 1 dB, respectively. These levels were selected based on pilot testing to yield high, intermediate, and low speech-recognition performance while avoiding floor and ceiling effects. The easy condition resulted in a mean performance of 91.61% of keywords repeated correctly (SD = 0.87), the medium condition in 76.08% (SD = 1.32), and the hard condition in 50.47% (SD = 2.43). Performance remained stable across the first and second halves of the experiment (see Figure S1).

The laboratory setup was calibrated using a Brüel & Kjær 2250 sound level meter and BCK Type 4231 calibrated sound source, such that background noise alone was played at a sound intensity of 65 dB LAeq. Overall sound levels of combined speech-in-noise stimuli varied according to the SNR.

To mitigate confounding effects of sentence content and difficulty, participants were assigned approximately equally across three mappings between sentences and difficulty levels, such that approximately one third of participants received each mapping. This ensured that each sentence occurred at each difficulty level across participants, preventing any given sentence from being systematically associated with a specific difficulty level. Reward levels were manipulated independently of this mapping and were fully crossed with difficulty levels.

In addition, sentence order was varied to control for order effects while maintaining a balanced distribution of difficulty and reward conditions across the experiment. One of four stimulus orders - the original order, an order with the first and second halves exchanged, a fully reversed order, or an order in which each half was reversed independently - was assigned pseudorandomly to each participant. All orders were constructed such that difficulty and reward levels were evenly distributed across the first and second halves of the experiment, while also varying across consecutive trials to avoid blocked presentation. A participant-specific Excel file containing this information was generated for stimulus presentation in PsychoPy.

#### Rest stimuli: City scenes in silence

*Rest* stimuli consisted of 26 different grey-scale images of city scenes from Copenhagen obtained from the VisitCopenhagen media platform (https://platform.crowdriff.com/m/visitcopenhagen), with photographs by Thomas Høyrup Christensen, Daniel Jensen, Daniel Rasmussen, and Martin Auchenberg. Images were assigned to the 126 trials in fixed, predefined order. The brightness of the images was normalized and adjusted to match the brightness of the previous screen to minimize light-induced changes in pupil dilation. When the participant chose to rest, the image assigned to that trial was displayed for 10 s in silence.

### Procedure

The experiment involved two test visits, scheduled six to ten days apart. The first test visit comprised an audiological check-up, fitting of the test hearing aids used in the experiment, including a program in which all additional noise-reduction features were disabled, and estimation of the individual difficulty levels using an SRT procedure. Cognitive assessments were also administered during this visit but are not analyzed in the present study.

The SRT estimation procedure was based on Keidser et al. (2013). Three rounds of SRT estimation were conducted, with a maximum of 30 sentences per round. The first round was considered a training round and served to familiarize the participant with the procedure: data from this round were not included in the SRT calculation. SRT estimation followed a sentence-based scoring procedure, with a success criterion of at least two of the three keywords repeated correctly.

The procedure started at an SNR of 10 dB. Correct responses result in a decrease of SNR, incorrect responses in an increase in SNR, converging on the 50% SRT. The procedure involves 3 phases: In phase 1, 5 dB steps were used until at least four sentences had been completed and at least one reversal had occurred. In phase 2, 2 dB steps were used until at least four additional sentences had been completed and the estimated standard error (SE) was ≤ 1.0 dB. In phase 3, 1 dB steps were used until at least 16 sentences had been completed from the start of phase 2 and the estimated SE was ≤ 0.8 dB. The SRT was calculated as the mean SNR across phases 2 and 3. Custom MATLAB scripts were used to present DAST stimuli in noise.

During the second visit, participants completed the listening-effort discounting paradigm, which constituted the main task, with concurrent fNIRS and pupillometry recordings. Before starting the main task, thorough instructions were given in oral and written form, and participants completed a familiarization phase and a pre-task. The pre-task, which lasted approximately 15 minutes, employed a broader set of effort-reward combinations (5 difficulty levels × 5 reward levels) than the main task (3 × 3) to characterize effort discounting prior to fatigue induction. Because the pre-task addressed research questions beyond the scope of the present study, it is not described further and its data were not included in the present analyses.

During familiarization, the experimenter guided participants through five example trials in which all difficulty levels were presented consecutively and paired with varying reward values. This allowed participants to become familiar with the listening demands associated with each difficulty level and with the subsequent fatigue-rating procedure. To ensure participants understood the offer visualization and response procedure, no time constraints were imposed. Subsequently, participants completed 10 trials in which difficulty and reward levels were randomly paired, using the same timing constraints as the pre-task and main task, to familiarize them with the experimental flow.

## Analysis and Results

### Data summary and descriptive statistics of fatigue ratings and decision behavior

#### Fatigue Ratings

Participant-level mean fatigue ratings ranged from 2.18 to 79.24, with an overall mean of 42.13 (SD=22.07;**Figure 4A**). Across participants, fatigue ratings increased over the course of the experiment, from a mean baseline rating of 15.43 (SD=17.25) to a mean final rating of 70.47 (SD= 24.32). However, individual fatigue trajectories showed considerable within-participant variability. To assess whether participants showed evidence of intermittent recovery, we quantified fatigue reversals, defined as changes in the direction of successive fatigue ratings. Fatigue ratings changed direction on average 10 times per participant (SD = 18.4), with unchanged ratings treated as a continuation of the preceding trend. On average, fatigue ratings increased by 0.41 units (SD = 4.0) from one trial to the next (Δ-fatigue).

**Figure 4.**
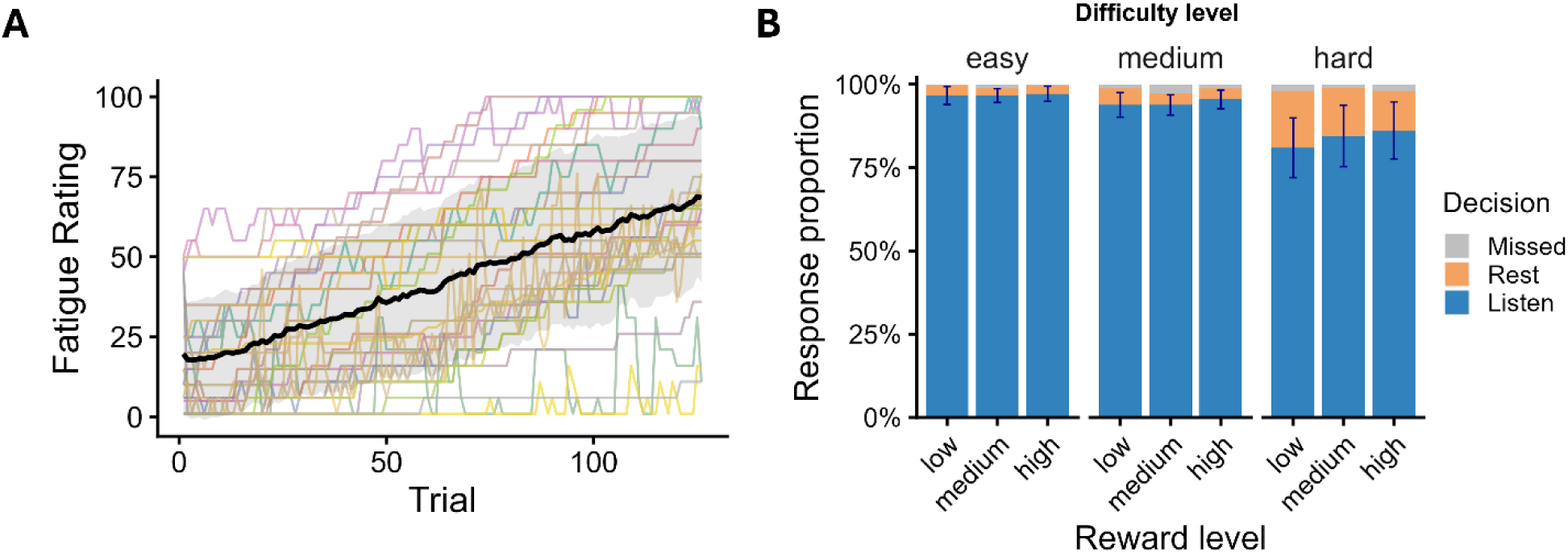
Fatigue ratings and listen-versus-rest decisions. **A.** Subjective fatigue ratings for each participant, obtained on every trial using a scale from 0 to 100. The black line represents mean ratings with one standard deviation shown in grey, illustrating an overall increase in self-reported fatigue over the course of the experiment. **B**. Observed response proportions for the nine reward-by-difficulty combinations, showing that participants generally chose to listen, with rest choices occurring primarily for high-difficulty offers. Listening, rest, and missed choices are shown separately. Error bars indicate S5% confidence intervals for the proportion of listening choices.

#### Listen-versus-Rest decisions

Participants accepted on average 91.7% of the 126 listening offers (SD= 9.3%), chose to rest on 7% of trials (SD=9.3%), and missed 1.3% of trials (SD=2.3%) (**Figure 4B** shows the mean decision proportions across reward and difficulty levels). Seven of the 30 participants never chose to rest during the experiment.

### Behavioral analysis and results

The behavioral analyses addressed whether listening-related fatigue accumulated over the course of the experiment, how moment-to-moment changes in fatigue were associated with task experiences, and whether participants’ willingness to exert effort for reward changed over time or with subjective fatigue.

Behavioral analyses included all 30 participants unless stated otherwise. Analyses were conducted in R using linear or generalized linear mixed-effects models, as appropriate. Categorical predictors were treatment-coded. Continuous trial-level predictors, including trial number, subjective fatigue ratings, and fatigue-change measures, were z-scored within participants; PTA4 was z-scored across participants.

Participant-specific random intercepts were initially considered for all models. However, models predicting changes in fatigue relative to the previous trial (Δ-fatigue) consistently resulted in singular fits, indicating negligible between-participant variance after accounting for the fixed effects. These analyses were therefore conducted using linear models without participant random intercepts.

For each of the fatigue and delta-fatigue models, a primary additive model was compared with a more complex model including theoretically motivated two-way interactions. Model selection was based on the Akaike Information Criterion (AIC). Models differing by less than two AIC units (ΔAIC < 2) were considered to have comparable support (Burnham and Anderson, 2004), in which case the more parsimonious model was retained for inference. Overall effects were assessed using ANOVA tests, and significant categorical predictors were followed by contrasts of estimated marginal means with Bonferroni correction within each contrast family..

#### Fatigue Ratings

##### Does subjective fatigue increase over time?

To assess whether subjective fatigue increased over the course of the experiment, fatigue ratings were modeled as a function of trial number and PTA4. The model confirmed that subjective fatigue increased over time (β = 15.43, *p* < .001; **Figure 5A**). PTA4 was not a significant predictor of fatigue ratings after accounting for participant random effects (β = 5.78, SE = 3.97, *p* = .16).

**Figure 5.**
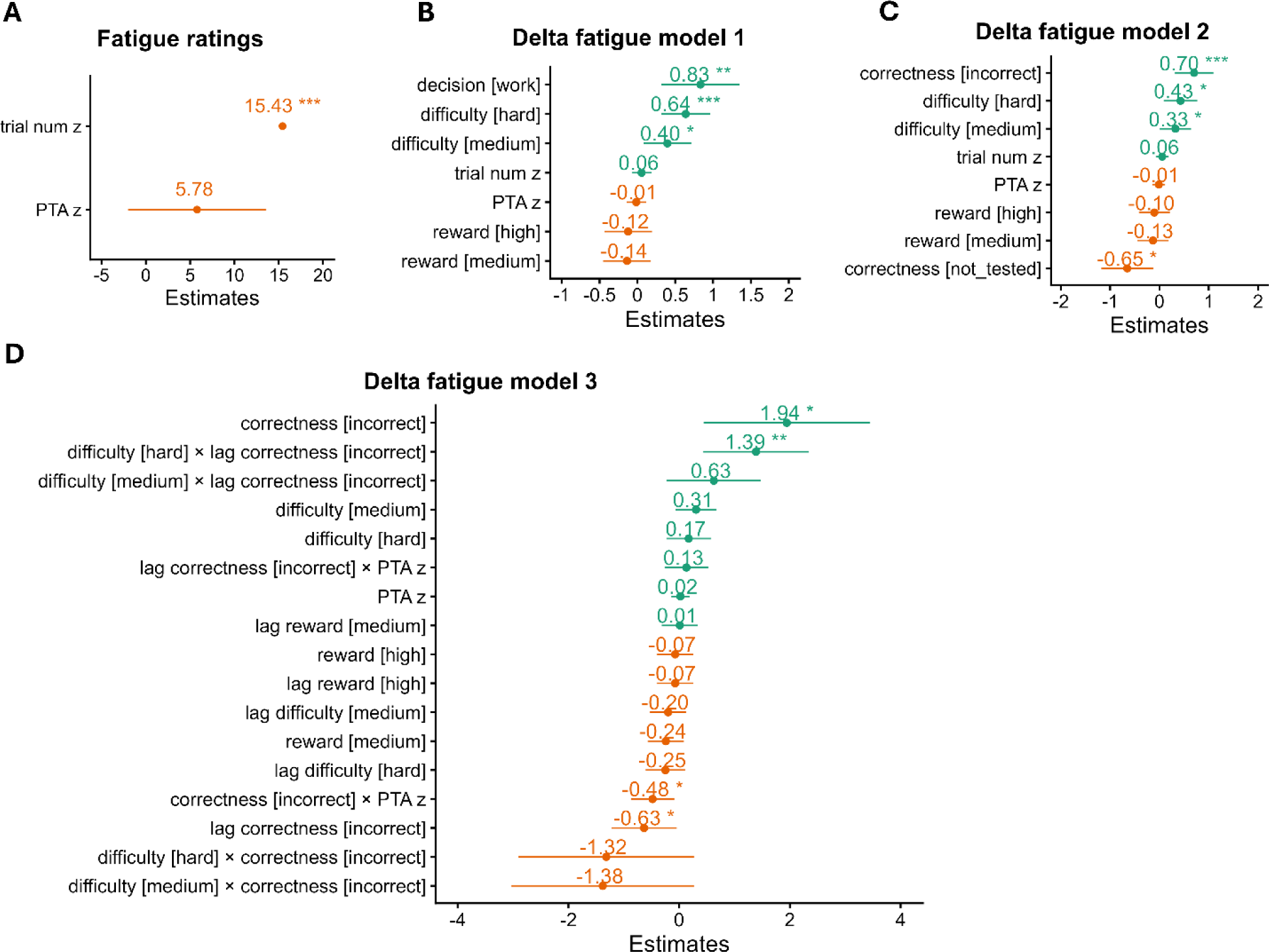
Fatigue model coefficients. **A.** Regression coefficients from a linear mixed effects model predicting absolute fatigue ratings from trial number and hearing loss severity (PTA4), demonstrating a significant increase of absolute ratings by trial number. **B.** Regression coefficients from a linear model predicting changes in fatigue ratings relative to the previous trial (Δ-fatigue) from listening difficulty, reward, decision (listen versus rest), trial number, and hearing loss severity (PTA4) **C.** Model coefficients from a linear model predicting changes in Δ -fatigue from the current trial success (=correctness), controlling for effects of difficulty and reward. **D.** Model coefficients from a linear model predicting changes in Δ -fatigue from the current-trial success, and previous-trial success, controlling for effects of difficulty and reward. This model was restricted to listening trials only. In all panels positive coefficients are plotted in green, and negative coefficients in orange. Significance is indicated by asterisks * p<0.05, ** p<0.01, *** p<0.001. Horizontal bars indicate S5% confidence intervals.

##### Which factors predict changes in fatigue ratings?

To examine which task-related factors were associated with changes in subjective fatigue relative to the previous trial, delta fatigue was modelled as a function of current-trial difficulty, reward, listening-versus-rest decision, trial number, and PTA4. An exploratory interaction model including all two-way interactions among these predictors reduced residual variance relative to the additive model but did not meaningfully improve overall model fit once model complexity was taken into account (ΔAIC = 0.06). The more parsimonious additive model was therefore retained for inference; details of the model comparison and the full exploratory interaction model are reported in Tables S1-S3.

The additive model revealed significant effects of task difficulty (F(2, 3723) = 7.78, p < .001) and decision (F(1, 3723) = 10.03, p = .0016), but not reward (F(2, 3723) = 0.44, p = .65), trial number (F(1, 3723) = 0.71, p = .40) or PTA4 (F(1, 3723) = 0.05, p = .82, model coefficients are shown in **Figure 5B**).

For decision, model results showed that fatigue ratings decreased following rest trials but increased following listening trials (estimated marginal means: rest = −0.37, listen = 0.47; contrast rest vs.listen: Δ = −0.83, SE = 0.26, p = .0016. This pattern is consistent with the raw data shown in **Figure 6A**, suggesting that rest choices were associated with relative reductions in fatigue.

**Figure 6.**
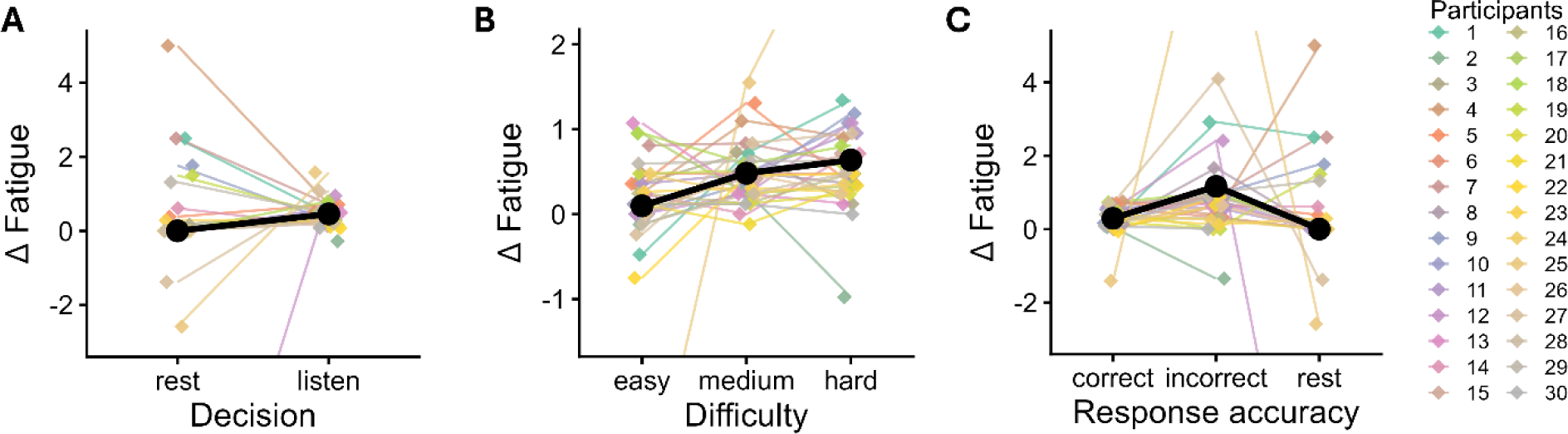
Δ-fatigue raw data. **A.** Δ-fatigue split by listen-versus-rest trials. On average (shown in black), participants increased their fatigue ratings more after listening than after resting. Note that one participant’s rest value fell outside the plotted y-axis range (Δ-fatigue = −12.0). **B.** Δ-fatigue split by difficulty level. On average (shown in black) participants increased their fatigue ratings more on medium or hard than on easy trials. Note that two values fell outside the plotted y-axis range (Δ-fatigue = −3.8 and 3.S). **C.** Δ-fatigue during listening trials split by trial success. On average (shown in black), participants increased their fatigue ratings more on incorrect than on correct trials (even though feedback about trial success had not yet been provided at the time of the rating). Note that two values fell outside the plotted y-axis range (Δ-fatigue = −12.0 and 10.4).

For difficulty, estimated marginal means indicated changes in fatigue ratings relative to the previous trial of −0.29, 0.10, and 0.34 units for easy, medium, and hard trials, respectively (the corresponding raw data are shown in **Figure 6B**). Pairwise comparisons indicated that fatigue increased less on easy trials than on medium trials, Δ = −0.40, p = .042, and hard trials, Δ = −0.64, p < .001. The difference between medium and hard trials was not significant, Δ = −0.24, p = .41.

Difficulty and reward were coded according to the characteristics of the offer presented to participants. Thus, difficulty and reward main effects reflect the consequences of being offered trials with different effort and incentive levels, irrespective of whether participants chose to complete or decline the trial. To distinguish whether effects of difficulty reflect merely being offered a more effortful listening condition or actually exerting effort, we also examined the Difficulty × Decision interaction in the more complex model. Although the interaction did not reach significance, *F*(2, 3704) = 2.75, *p* = .064, estimated marginal means indicated numerically higher fatigue following listening than rest decisions for medium and hard difficulty offers, whereas little variation was observed across difficulty levels when participants chose to rest. This pattern is consistent with, but does not provide strong evidence for, a greater role of effort exertion than offered task difficulty per se.

For reward, the distinction between offered and received reward is more closely tied to trial success than to the decision to listen versus rest, because reward points could only be obtained following a correct response. We therefore examine this question in the following analysis.

##### Does trial success predict changes in fatigue ratings?

To distinguish the effects of choosing to listen or rest from the effects of successful versus unsuccessful task performance on delta fatigue, a second delta-fatigue model was fitted in which the decision-predictor was replaced by a trial success predictor with three levels: correct listening trials, incorrect listening trials, and rest trials (‘not tested’). Because fatigue was rated before feedback and reward presentation, trial success reflects participants’ perceived performance at the time of rating. The model included trial success, trial number, difficulty, reward, and PTA4 as fixed effects.

An exploratory interaction model including all two-way interactions among trial number, difficulty, reward, trial success, and PTA4 did not improve model fit once model complexity was taken into account (ΔAIC = 5.08), favoring the additive model). The more parsimonious additive model was therefore retained for inference; details of the full exploratory interaction model are reported in Tables S4-S5.

The additive model showed significant effects of task difficulty, *F*(2, 3722) = 3.42, *p* = .033, and trial success, *F*(2, 3722) = 11.27, *p* < .001. There were no significant effects of trial number, *F*(1, 3722) = 0.73, *p* = .392, reward, *F*(2, 3722) = 0.37, *p* = .693, or PTA4, *F*(1, 3722) = 0.04, *p* = .846. Model coefficients are shown in **Figure 5C**.

Estimated marginal means indicated significant differences between all levels of trial success. Fatigue increased more on incorrect listening trials than on correct listening trials (Δ = 0.70, SE = 0.20, *t*(3722) = 3.54, *p* = .001), consistent with the raw data shown in **Figure 6C**. Increases were also larger on incorrect trials than on rest trials (Δ = 1.36, SE = 0.30, *t*(3722) = 4.49, *p* < .001) and on correct listening trials than rest trials (Δ = 0.65, SE = 0.27, *t*(3722) = 2.43, *p* = .045). For task difficulty, fatigue increases were larger following hard than easy trials (Δ = 0.43, SE = 0.17, *t*(3722) = 2.46, *p* = .042), whereas the other pairwise comparisons were not significant after Bonferroni correction.

Visual inspection of the raw data suggested that one participant had a particularly strong influence on the effect. A leave-one-out analysis confirmed that this participant contributed more strongly than others; however, the effect of trial success remained significant when this participant was removed from the analysis (*F*(2, 3597) = 4.63, *p* = .010)

Given that reward reflected the incentive offered on a given trial rather than the reward ultimately obtained, we examined the reward × trial success interaction from the complex model. Consistent with the absence of a main effect of reward, the results provided no evidence that the association between reward level and changes in fatigue depended on trial success (*p* = .079; Table S4).

##### Do trial success and feedback predict changes in fatigue ratings?

To examine effects of trial success more specifically, a third model predicting delta fatigue was fitted to test whether changes in fatigue were more strongly associated with current-trial success (reflecting participants’ perceived performance before feedback) or with feedback from the previous trial. This analysis was restricted to listening trials that were preceded by another listening trial (3,215 trials across 30 participants), ensuring that both current-and previous-trial success were defined and could be disentangled from listen-versus-rest decisions.

The model included current trial success, previous-trial success, while controlling for current-and previous-trial difficulty, current-and previous-trial reward, and PTA4 as fixed effects. The additive model of these predictors was compared with a more complex model that additionally included theoretically motivated interactions involving current-and previous-trial success. Because the interaction model showed improved fit relative to the additive model (ΔAIC = 4.80), the interaction model was retained for inference. Details of the additive model are reported in Tables S6-S7.

Type III tests revealed significant effects of current-trial success, *F*(1, 3197) = 6.44, *p* = .011, and previous-trial success, *F*(1, 3197) = 4.43, *p* = .035. In addition, significant interactions were observed between current-trial success and PTA4, *F*(1, 3197) = 5.77, *p* = .016, and between current-trial difficulty and previous-trial success, *F*(2, 3197) = 4.13, *p* = .016. Model coefficients are shown in **Figure 5D**.

Follow-up analyses indicated that the association between PTA4 and delta fatigue differed between correct and incorrect trials. For correct trials, PTA4 was not associated with delta fatigue (slope = 0.09, SE = 0.10, 95% CI [-0.12, 0.29]), whereas for incorrect trials higher PTA4 values were associated with smaller increases in delta fatigue (slope = −0.39, SE = 0.20, 95% CI [-0.78, −0.005]). Thus, the difference in delta fatigue between correct and incorrect trials decreased with increasing PTA4.

The association between previous-trial success and subsequent changes in fatigue depended on the difficulty of the current trial. On easy trials, delta fatigue was higher when the preceding trial had been correct rather than incorrect (1.15 vs. 0.52; Δ = 0.64, *p* = .034). No such difference was observed for medium-difficulty trials (0.77 vs. 0.76; Δ = 0.01, *p* = .97). On hard trials, the pattern tended to reverse, with higher delta fatigue following an incorrect than a correct previous trial (1.42 vs. 0.67; Δ = −0.75, *p* = .053).

Taken together, these findings suggest that changes in fatigue were associated not only with task difficulty but also with performance-related factors from the current and preceding trial. Thus, the effect of difficulty observed in the initial Δ-fatigue model reflected an overall effect across listening and rest trials, whereas the third Δ-fatigue model, restricted to consecutive listening trials, indicated that the relationship between difficulty and fatigue also depended on previous-trial success.

#### Listen-versus-rest decisions

##### Which factors predict listen-versus-rest decisions?

Fatigue ratings were highly correlated with trial number (r = .93, 95% CI [.92, .93], p < .001), with trial number accounting for approximately 86% of the variance in fatigue ratings (R^2^ = .86). To separate fatigue-related effects from general time-on-task effects, fatigue was orthogonalized with respect to trial number by regressing fatigue ratings on trial number and retaining the residuals. To examine predictors of listen-versus-rest decisions, we fitted a binomial generalized linear mixed model including task difficulty, reward, trial number, previous-trial success, previous-trial change in fatigue, residualized fatigue ratings, PTA4, final fatigue ratings, and the number of fatigue reversals as fixed effects, with participant included as a random intercept. Fatigue reversals, defined as changes in the direction of successive fatigue ratings, were included to capture individual differences in the extent to which fatigue trajectories exhibited short-term decreases and recoveries rather than a monotonic increase over time. Previous trial success was coded at three levels in this model (correct or incorrect for listening trials and ‘not tested’ when the previous trial was a rest trial, as in Δ-fatigue model 2. Participants who never chose to rest during the experiment (n = 7) were excluded because their data provided no information for estimating predictors of listen-versus-rest choices.

Type-III Wald tests revealed significant effects of task difficulty (χ^2^(2) = 167.16, p < .001), reward (χ^2^(2) = 6.79, p = .03), trial number (χ^2^(1) = 40.94, p < .001), previous-trial success (χ^2^(2) = 9.24, p < .001) and fatigue reversals (χ^2^(1) = 5.32, p = 0.02) on the probability of accepting offers. No significant effects were observed for residualized fatigue ratings, previous-trial delta fatigue, PTA4 or final fatigue ratings (coefficients of this model are shown in **Figure 7B**).

**Figure 7.**
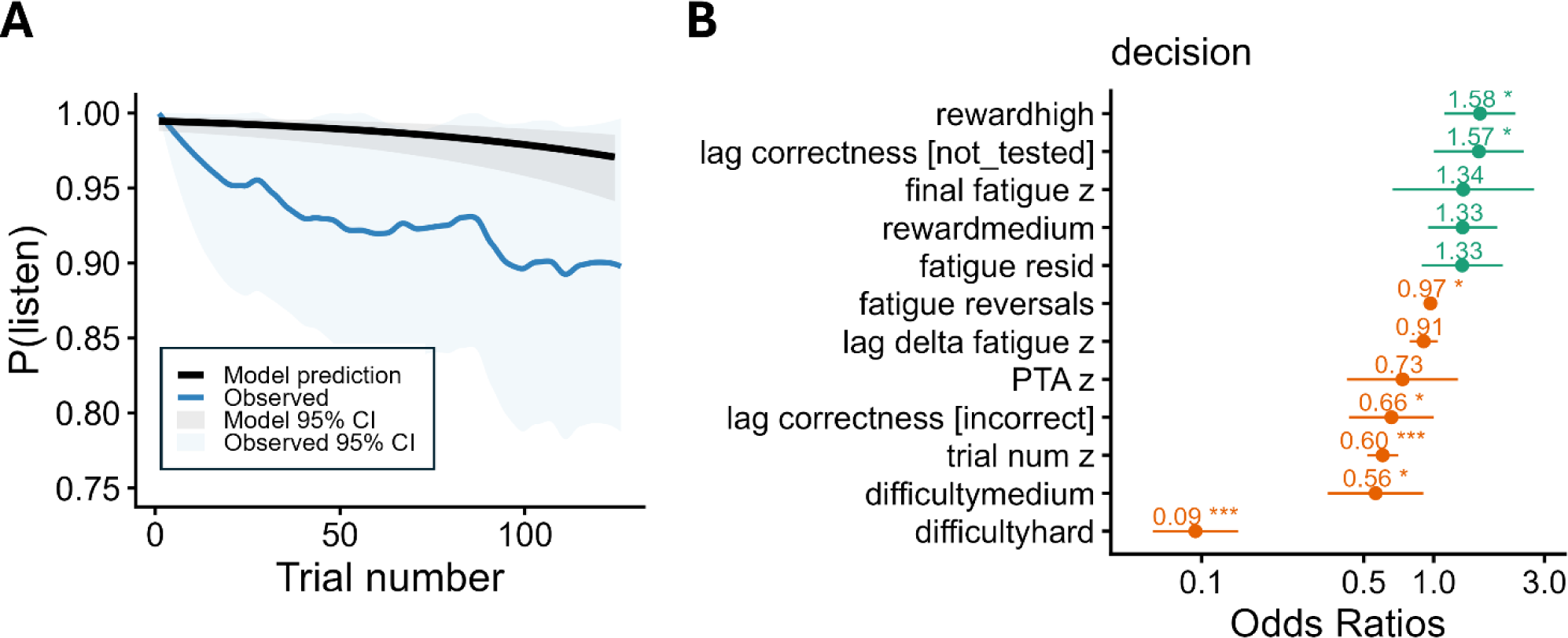
Predictors of listen-versus-rest decisions. **A.** Observed and model-predicted probabilities of choosing to listen (P(listen)) as a function of trial number. The blue line shows a smoothed estimate of the observed proportion of listening choices across trials, with corresponding S5% confidence intervals shown in light blue. The black line shows probabilities predicted by the mixed-effects model, with all other predictors held constant showing that participants became increasingly likely to choose rest over the course of the experiment, although listening choices remained near ceiling. The predicted trajectory reffects the isolated effect of trial number and does not fully reproduce the observed choice pattern which was also affected by other factors than trial number (see B). **B.** Odds ratios and S5% confidence intervals from the mixed-effects model predicting listen-versus-rest decisions. Higher task difficulty, later trial numbers, and a greater number of fatigue reversals were associated with reduced odds of choosing to listen, whereas higher rewards increased the odds of choosing to listen. Participants were also more likely to choose to listen following a rest trial than following an incorrect listening trial. Horizontal bars indicate S5% confidence intervals and asterisks statistical significance: * p<0.05, ** p<0.01, *** p<0.001.

Estimated probabilities of choosing to listen were 98.7%, 97.7%, and 87.5% for easy, medium, and hard trials, respectively. Pairwise comparisons indicated significantly higher odds of choosing to listen for easy than hard trials (OR = 10.62, p < .0001) and for medium than hard trials (OR = 5.97, p < .0001).

Estimated probabilities of choosing to listen were 95.6%, 96.7%, and 97.2% for low, medium, and high reward trials, respectively. Pairwise comparisons indicated significantly lower odds of choosing to listen for low-than high-reward offers (OR = 0.63, p = .033). Contrasts involving medium reward were not significant (OR = 0.75, p = .31; OR = 0.84, p = 1.00). **Figure 4B** shows the corresponding raw data across reward and difficulty levels.

Trial number was associated with reduced odds of choosing to listen (β = −0.51, OR = 0.60, χ^2^(1) = 40.94, p < .001), indicating that participants became more likely to chose rest over the course of the experiment (**Figure 7A)**. However, because listening choices remained near ceiling, the corresponding change in predicted probabilities was small, decreasing from 99.5% on the first trial to 97.0 % on the last trial.

Estimated probabilities of choosing to listen were 96.5%, 94.8%, and 97.7% following correct, incorrect, and rest trials, respectively. Pairwise comparisons revealed significantly higher odds of choosing to listen following rest than following incorrect trials (OR = 2.38, p = .007). In contrast, the differences between correct and incorrect trials (OR = 1.52, p = .15) and between correct and rest trials (OR = 0.64, p = .15) did not reach significance after Bonferroni correction. These results suggest that the overall effect of previous trial success was primarily driven by the contrast between rest and incorrect trials rather than by differences between correct and incorrect listening trials.

A greater number of fatigue reversals, reflecting a more fluctuating fatigue profile, was associated with reduced odds of choosing to listen (β = −0.032, OR = 0.97, χ^2^(1) = 5.32, p = .021), suggesting that participants with more variable fatigue trajectories tended to choose rest more often.

We expected that as time on task and fatigue increased, difficult trials with small rewards would be increasingly avoided. To test this hypothesis, we fitted two follow-up GLMMs predicting listen-versus-rest decisions as a function of difficulty × reward × trial number and difficulty × reward × fatigue interactions, respectively, with participant included as a random intercept.

Neither model revealed significant interaction effects (Tables S8-S9), indicating that increasing fatigue and time on task did not alter how participants weighed reward and task difficulty when deciding whether to listen or rest. Thus, we found no evidence that the effects of difficulty and reward on listening decisions changed as a function of either time on task or subjective fatigue.

##### Are listen-versus-rest decisions better predicted by trial number or fatigue ratings?

To examine whether listen-versus-rest decisions were better accounted for by time on task or by subjective fatigue, two additional models were estimated and compared using AIC: one including trial number only and one including fatigue ratings only. Again, participants with no variability in their choices (i.e., those who never chose to rest) were excluded from this analysis.

Both models revealed significant effects of their respective predictors on listen-versus-rest decisions (trial number: χ^2^(1) = 34.94, p < .001; fatigue ratings: χ^2^(1) = 28.29, p < .001). However, model comparison indicated that the model including trial number provided a better fit than the model including fatigue ratings (ΔAIC = 7.3), suggesting that time on task captured decision-relevant variance beyond that explained by subjective fatigue ratings alone.

### fNIRS findings

The behavioral analyses indicated that subjective fatigue accumulated over the course of the experiment and that participants became less likely to accept listening offers over time. However, there was no evidence that fatigue or time on task altered how reward and difficulty were weighted during decision making. In line with these behavioral findings, we hypothesized that frontal cortical oxygenation would increase as fatigue accumulated over the course of the experiment, reflecting increased neural engagement during the evaluation of whether effortful listening was worth the potential reward. We first examined whether neural responses to reward and difficulty during accepted offers differed between the first and second halves of the experiment (Analysis 1). This analysis provided a simplified assessment of time-on-task effects and allowed us to explore whether the neural processing of reward and difficulty in accepted offers changed over time. To further characterize any observed half-related effects, we subsequently assessed whether frontal cortical activation varied as a function of continuous measures of time on task and subjective fatigue (Analysis 2).

All fNIRS analyses were restricted to accepted offers (listening trials). Although cortical activity was analyzed during offer evaluation, listening and rest trials differed in their subsequent auditory stimulation, response requirements and temporal structure. Restricting analyses to listening trials therefore avoided potential confounds related to trial type.

fNIRS data preprocessing and first-level analyses were conducted in Python 3.12.5 using MNE-Python (v1.8.0; Gramfort et al., 2013) and MNE-NIRS (v0.7.1; Luke, Larson, et al., 2021), with additional use of nilearn (Abraham et al., 2014) and statsmodels (Seabold & Perktold, 2010) packages. Second-level analysis were conducted in R unless stated otherwise. In line with common practices for fNIRS analyses (Yücel et al., 2021, 2025), interpretations were based on oxygenated hemoglobin (HbO) results, given its higher signal-to-noise ratio and sensitivity to task-related changes. Deoxygenated hemoglobin (HbR) signals are shown for completeness but were not included in the statistical analyses. HbO concentration changes are reported in micromolar units (µM, equivalent to µmol/L).

#### Preprocessing

Raw light intensity data were converted to optical density, followed by the rejection of channels expected to poorly index neural responses. Channels with scalp coupling index (SCI) smaller than 0.77, or a peak power below 0.2 in more than 30% of 10-s windows, or a coefficient of variation greater than 0.15 were excluded from further analyses (see also Pollonini et al., 2016). Datasets with more than 50% of all long channels marked as bad based on these criteria were excluded from further analysis. This was the case for four participants. For the remaining sample of N = 26 participants, on average 70.6 % (SD = 27.7 %) of all long channels were retained for analysis.

Temporal derivative distribution repair (Fishburn et al., 2019) was applied to reduce potential motion artefacts, before the data were converted to hemoglobin concentration changes using the modified Beer-Lambert law with a partial pathlength factor of 0.1. Hemoglobin data was then low-pass filtered with a 5^th^-order IIR Butterworth filter using a cut-off frequency of 0.5 Hz and subsequently downsampled to 2 Hz. Finally, short-channel regression was performed in a separate GLM, in which HbO and HbR signals from all eight short channels were included as nuisance regressors to account for extracerebral physiological activity. The weighted contribution of the short-channel regressors was then reconstructed and subtracted from the original signal, yielding a corrected time series with reduced contamination from extracerebral signals. For one participant in whom no short channels of sufficient quality remained after bad-channel rejection, principal component analysis (PCA) was applied to the full dataset, and the first principal component was subtracted to account for global physiological noise.

#### Finite Impulse Response – General Linear Model

Predefined hemodynamic response functions, such as the SPM canonical (double-gamma) or the Glover HRF, commonly used in fMRI and fNIRS models, impose a fixed response shape derived from a limited set of stimuli, including auditory and sensorimotor responses (Glover, 1999). However, HRF morphology varies across regions (Prokopiou et al., 2022), motivating a more flexible approach, particularly for frontal regions of interest. We therefore used a finite impulse response (FIR) model, which allowed the HRF shape to be estimated without imposing a predefined temporal profile.

As the experiment served multiple purposes beyond fNIRS acquisition, including the investigation of decision-making behavior and additional physiological measures (not analyzed here), the paradigm was not specifically designed to isolate a hemodynamic response to the offer event. Consequently, each trial comprised a sequence of cognitive and sensory processes, with subsequent speech-in-noise perception and verbal repetition following the offer evaluation. Because these processes occurred in close temporal succession and their associated hemodynamic responses were expected to overlap, we modelled 23 s of each trial using an FIR approach. This allowed us to test whether task conditions, trial number, or fatigue preferentially modulated early delay bins, which were expected to reflect offer evaluation most strongly, rather than later bins that could also reflect speech perception and verbal responses (see **Figure 11** for an illustration of the FIR modelling approach used in Analysis 2).

#### Analysis 1 – Is frontal cortical activation in relation to offer evaluation modulated by task conditions and first versus second experimental half?

To analyze whether frontal cortical activation during evaluation of accepted offers differed as a function of task conditions and experimental half within participants, first level design matrices were constructed using the *mne_nirs.experimental_design* module. FIR regressors spanned 0 to *23* s following stimulus onset, resulting in 46 delay bins at a sampling rate of 2 Hz. For each of the 18 task conditions (3 difficulty levels x 3 reward levels x first vs. second experimental half), separate FIR regressors were included, resulting in a total of 828 regressors. In addition, low-frequency drifts were modeled using cosine basis functions with a high-pass cutoff of 0.01 Hz. GLM estimation was performed on all long channels (excluding channels marked as bad) using an AR(5) noise model to account for temporal autocorrelation. Given that FIR estimates are susceptible to edge instabilities, typically resulting in boundary constraints that force responses to decay toward zero (Goutte et al., 2000), the first and last two bins are displayed for completeness but were not interpreted.

To quantify group effects of offer-related activity, HbO beta estimates were averaged within each participant across an early time window of 2-6 s following trial onset (bins 4-12), corresponding to the expected peak of the hemodynamic response to offer evaluation based on a canonical HRF. A linear mixed-effects model was fitted to data from the frontal ROI, with reward, difficulty, and half as fixed effects including all interactions. Fatigability, quantified as the z-scored slope of a linear regression fitted to each participant’s fatigue ratings over time, was included as a covariate intended to account for systematic variability in frontal activation beyond baseline differences captured by participant random intercepts. Variability across fNIRS channels within the frontal ROI was additionally accounted for by including channel as a random intercept in a crossed random-effects structure with participant. Significant effects were followed by planned contrasts of estimated marginal means.

## Results

In the early time window, frontal cortex activation differed as a function of difficulty, *F*(2, 9696.6) = 5.15, *p* = .006, and experimental half, *F*(1, 9696.3) = 55.94, *p* < .001. No main effects of reward or fatigability were observed. Importantly, half-related changes in activation depended on task condition, as indicated by significant reward × half, *F*(2, 9695.9) = 3.78, *p* = .023, difficulty × half, *F*(2, 9696.2) = 34.91, *p* < .001, and reward × difficulty × half interactions, *F*(4, 9695.9) = 8.73, *p* < .001. Model-estimated hemodynamic responses across all delay bins, shown separately for the frontal and temporal ROIs and for the first and second experimental halves, are represented in **Figure 8**. Full ANOVA results are reported in Table S10, and model coefficients are visualized in **Figure 9B**.

**Figure 8.**
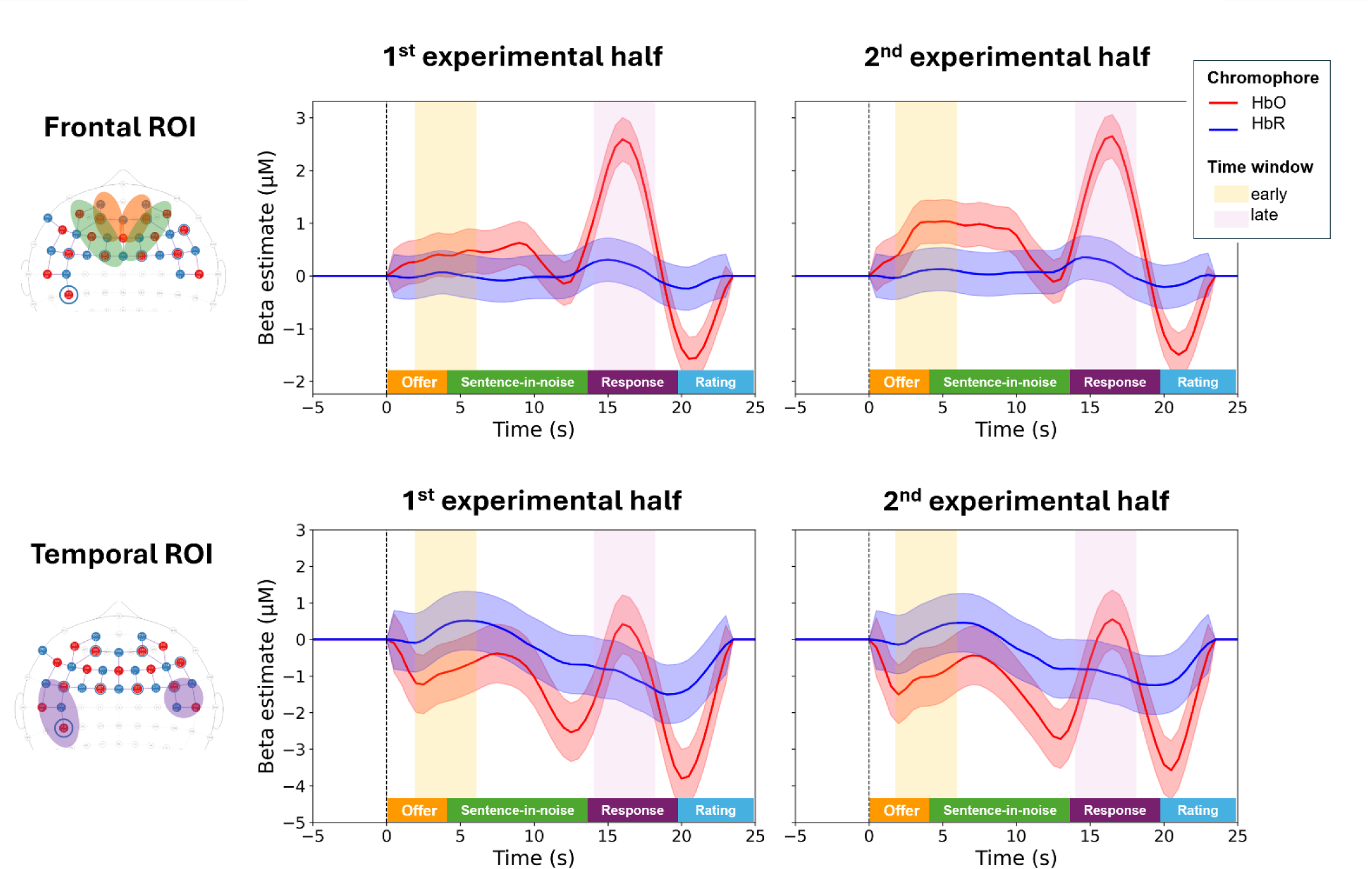
Model-estimated hemodynamic response functions across experimental halves. Group-level estimates of HbO and HbR concentration changes are shown in red and blue, respectively, for each ROI and experimental half.The visualization of the hemodynamic response across the full trial was based on one linear mixed-effects models for each ROI of the form: Beta ∼ −1 + Bin × Chroma × Half + (1|participant) + (1|channel). Note that, in the model used for statistical inference, delay bins were not retained as separate fixed effects. Instead, values were averaged across bins within participants and within the predefined time window, acknowledging that temporally adjacent bins are likely to be autocorrelated.

**Figure 9.**
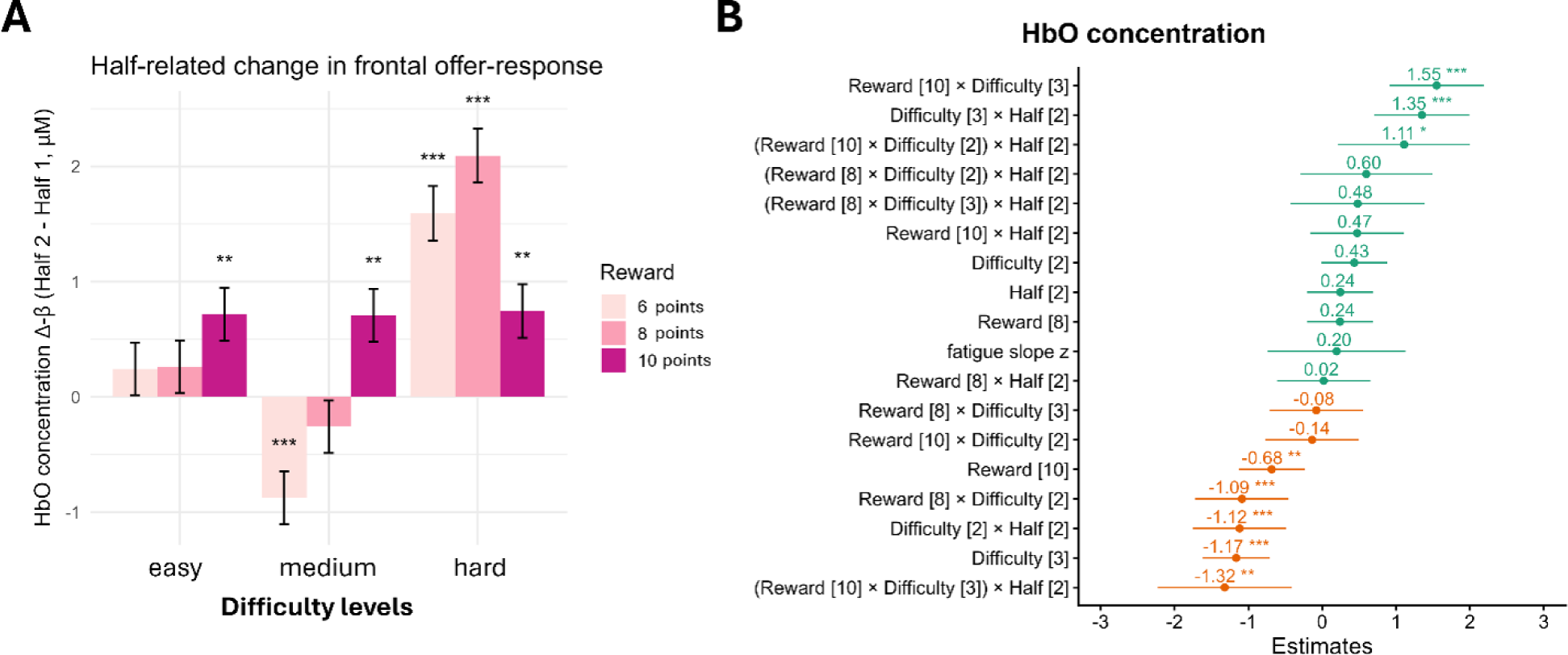
fNIRS analysis 1 – group-level results. **A**. Model-estimated change in frontal HbO response from the first to the second experimental half in the early offer-evaluation time window, shown separately for each difficulty and reward condition. Results show larger half-related increases for hard trials, particularly when hard trials were paired with low or medium rewards. Positive values indicate larger HbO responses in the second than in the first half. Asterisks indicate Bonferroni-corrected follow-up contrasts comparing the first and second half within each reward × difficulty condition. Error bars indicate standard errors of these contrasts. **B.** Fixed-effect coefficient estimates from the mixed-effects model predicting frontal HbO concentration in the early offer-evaluation time window. The model included reward, difficulty, experimental half, their interactions, and fatigue slope as fixed effects, with random intercepts for participant and channel. Horizontal bars indicate S5% confidence intervals. Positive coefficients are plotted in green, and negative coefficients in orange. Significance is indicated by asterisks * p<0.05, ** p<0.01, *** p<0.001. Horizontal bars indicate S5% confidence intervals.

Frontal oxygenation during offer evaluation in the early time window increased from the first to the second half of the experiment. Estimated marginal means indicated an increase from 0.35 µM, 95% CI [-0.48, 1.18], to 0.93 µM, 95% CI [0.10, 1.76], corresponding to a half-related increase of Δ = 0.51 ± 0.08 µM, *p* < .001.

Follow-up Bonferroni-corrected pairwise comparisons indicated lower HbO concentrations for medium-difficulty than easy trials (*p* = .004), whereas no other difficulty contrasts reached significance (estimated marginal means: easy = 0.80 µM, 95% CI [-0.04, 1.63]; medium = 0.49 µM, 95% CI [-0.34, 1.33]; hard = 0.63 µM, 95% CI [-0.2, 1.47], see **Figure 10** for participant-level first-level estimates across difficulty levels and experimental halves).

**Figure 10.**
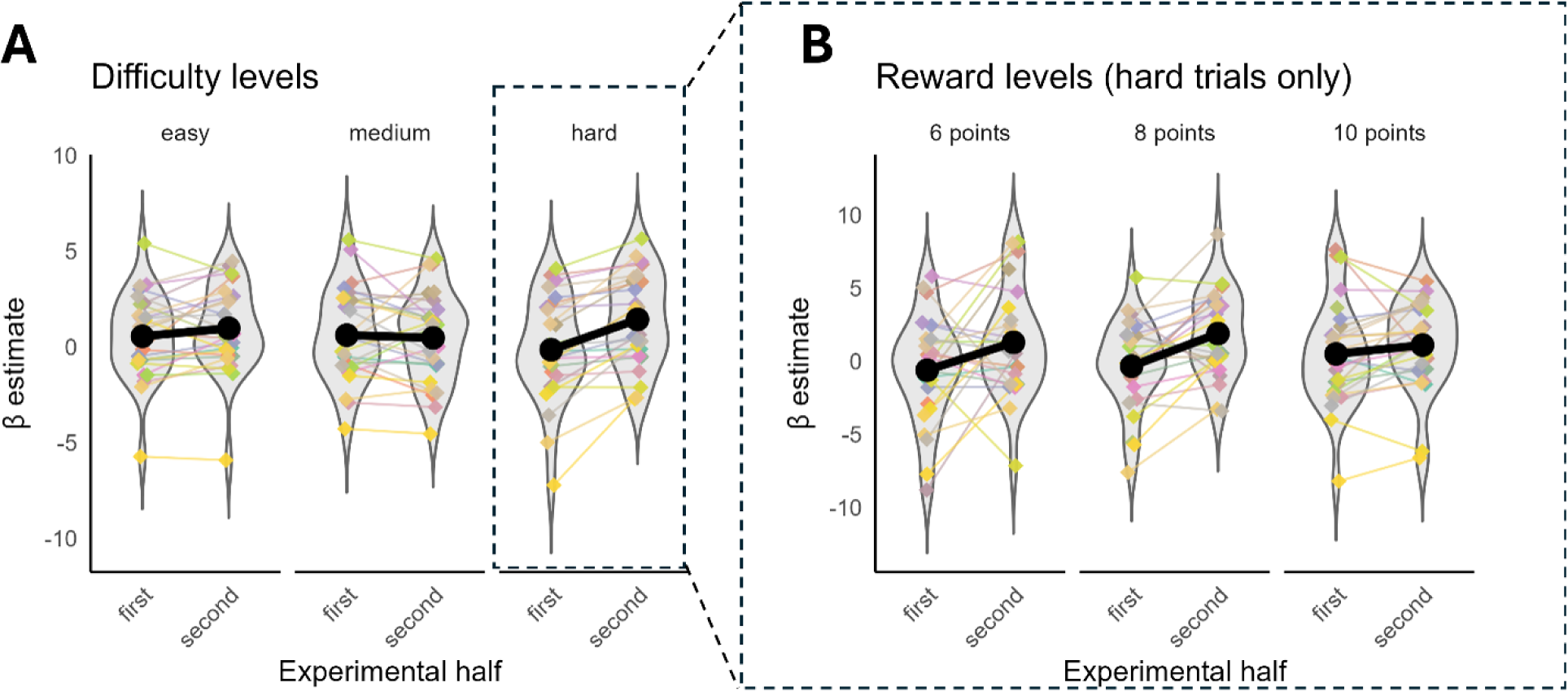
fNIRS analysis 1 – participant-level results. First-level HbO beta estimates of frontal oxygenation during the early time window are displayed. Colored points show individual participant estimates, grey density shapes indicate the distribution, and black lines indicate group means. **A.** Results split by difficulty and experimental half suggesting that mean frontal activation increases from the first to second experimental half are predominantly driven by difficult trials. **B.** Results for the hardest difficulty level, split by rewards and experimental half showing that increases in mean frontal activation within hard trials are predominantly driven by easy and medium reward conditions.

Follow-up Bonferroni-corrected contrasts showed that half-related changes in activation varied across reward and difficulty combinations (**Figure 9A)**. The largest increases from the first to the second half were found for high-difficulty offers paired with low rewards (Δ = 1.51 µM, *p* < .001) or medium rewards (Δ = 2.09 µM, *p* < .001).

To formally assess whether half-related changes differed across regions and time windows, an additional model including ROI, time window, and experimental half was fitted and predefined contrasts of interest performed. This model tested whether half-related changes differed between frontal and temporal ROIs and between the early offer-evaluation window and a later time window from 14 to 18 s, corresponding approximately to the verbal-repetition period (bins 28-36).

Significant main effects of window, *F*(1, 26797.8) = 110.96, *p* < .001, ROI, *F*(1, 30.7) = 44.69, *p* < .001, and half, *F*(1, 26798.3) = 5.69, *p* = .017, were observed, alongside interactions between window and ROI, *F*(1, 26797.8) = 23.92, *p* < .001 and half and ROI *F*(1, 26797.8) = 4.44, *p* = .035. The three-way half × ROI × window interaction approached but did not reach statistical significance, *F*(1, 26797.8) = 3.79, *p* = .052 (see Table S11 for full ANOVA results). Follow-up contrasts revealed that activation increased significantly from the first to the second half of the experiment only in the early frontal time window (Δ = 0.57 µM, SE = 0.11, *p* < .001), whereas no significant half-related effects were observed in the temporal ROI or late time window (all *p*s > .34, Table S12; see **Figure 8**).

Lastly, a separate control model was fitted to the early time window using data acquired from short channels to assess whether corresponding effects were present in extracerebral hemodynamic signals.

No main effect of half, *F*(1, 2809.3) = 0.33, *p* = .568, and no interactions involving half were observed (Table S13), suggesting that the frontal long-channel effects were unlikely to be driven by extracerebral hemodynamic changes. A main effect of reward was observed, *F*(2, 2809.0) = 3.43, *p* = .033, reflecting higher short-channel activity for reward 8 than reward 6 (*p* = .039). However, neither the reward × half nor the reward × difficulty × half interactions were significant, indicating that the condition-dependent half-related changes observed in the long-channel data were not mirrored in the short-channel signals (full ANOVA results are reported in Table S13).

### Analysis 2-Can half-related effects be further explained by linear time on task or subjective fatigue ratings?

Given the significant half-related increase in frontal cortical activation observed in Analysis 1, we conducted a follow-up analysis to determine whether this effect reflected a gradual linear increase in trial-by-trial response amplitudes over the course of the experiment or was associated with participants’ subjective fatigue ratings. In this analysis, reward and difficulty levels were not modelled separately. Instead, offer onsets from all accepted trials were combined and modelled as a single event type ‘offer’. Unlike Analysis 1, which summarized first-level results across multiple FIR bins within a predefined early time window, this analysis assessed whether trial-by-trial response amplitudes within each FIR delay bin were modulated by time on task or by subjective fatigue ratings.

The analysis was restricted to the frontal ROI and the first 13 seconds after offer onset (bins 1-26), excluding later parts of the trial that were less directly related to offer evaluation and had not shown half-related effects in Analysis 1. Consistent with the handling of FIR edge effects in Analysis 1, no conclusions were drawn from the first or last two bins.

Two first-level FIR-based GLMs were estimated to quantify HbO and HbR beta values for two sets of regressors: (1) FIR regressors capturing the mean evoked response to offer presentation (constant amplitude across trials) for each bin, and (2) modulation regressors capturing trial-by-trial variation in response amplitude in each bin. This separation allowed trial-by-trial variation in response amplitude to be estimated independently of the mean evoked response. In the first model, modulation regressors tested for systematic linear changes in response amplitudes across trial number (i.e., time on task; **Figure 11**). In the second model, modulation regressors tested whether trial-by-trial variations in response amplitudes were associated with subjective fatigue ratings. Low-frequency drifts were modelled using cosine basis functions (high-pass cutoff = 0.01 Hz), and beta values were estimated for all long channels (excluding channels marked as bad) using an AR(5) noise model.

**Figure 11.**
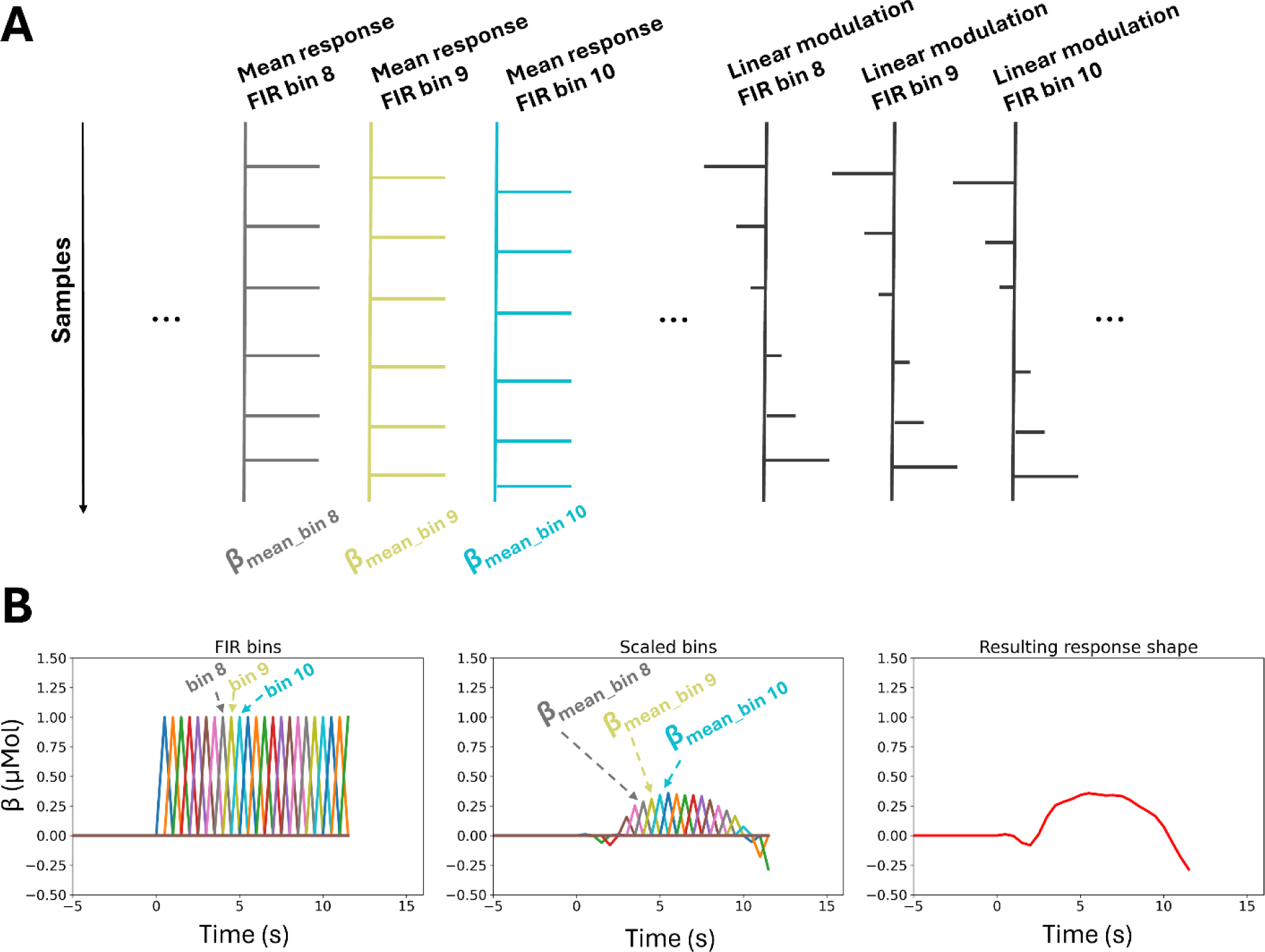
FIR design matrix for fNIRS analysis 2. **A** Schematic representation of an FIR design matrix with constant-amplitude and linearly modulated regressors. The example is illustrative only and does not depict data from the reported analysis. One set of regressors estimated the mean response amplitude in each FIR bin (left), while a second set of regressors modelled trial-by-trial modulation of response amplitudes within each bin (right), testing whether response amplitudes changed linearly across trials. In a second analysis (not shown), these modulation regressors were scaled by the individual z-scored subjective fatigue ratings rather than trial number. **B.** Example of a response estimated from 22 FIR bins at 2 Hz for a single regressor type (mean response). Left: FIR regressors corresponding to the design matrix. Middle: FIR regressors scaled by their estimated beta coefficients following GLM fitting. Right: reconstructed fNIRS response obtained from the beta estimates across FIR bins. For simplicity, the corresponding illustration of the second regressor type shown in panel A (linear modulation), which captures linear changes in response amplitude across trials within each FIR bin, is not shown here.

For each FIR-GLM, group-level inference was performed using linear mixed-effects models implemented in the Python package *statsmodels*. For each model, fixed effects included FIR delay bin, chroma (HbO versus HbR), and regressor type (mean response versus modulation effect), including all interactions. Random intercepts were included for participants and channels in a crossed random-effects structure to account for repeated observations within participants and variability across channels. Resulting *p*-values for fixed effects were corrected for multiple comparisons using false-discovery-rate (FDR) correction. Based on the observed time-on-task modulation effects, the model was repeated for the temporal ROI. Given that Analysis 1 indicated stronger modulation in the frontal than in the temporal ROI, we hypothesized that no significant linear modulation effects would be observed in the temporal ROI.

## Results

Two models testing delay-bin-specific modulations of the hemodynamic response were fitted, separating the average response amplitude from trial-by-trial modulation effects. Across both models, the mean offer response showed significant HbO values in delay bins 3-23. HbO values increased over the first 9 seconds and then decayed within the modelled time window, while HbR showed no significant deviation from zero (**Figure 12A** and **B** upper panels). The time-on-task modulation analysis revealed significant linear increases in response amplitudes for delay bins 6-11 and 14-24, with the strongest effect observed at delay bin 9 (4.5 s after offer onset; **Figure 12A**, lower panel).

**Figure 12.**
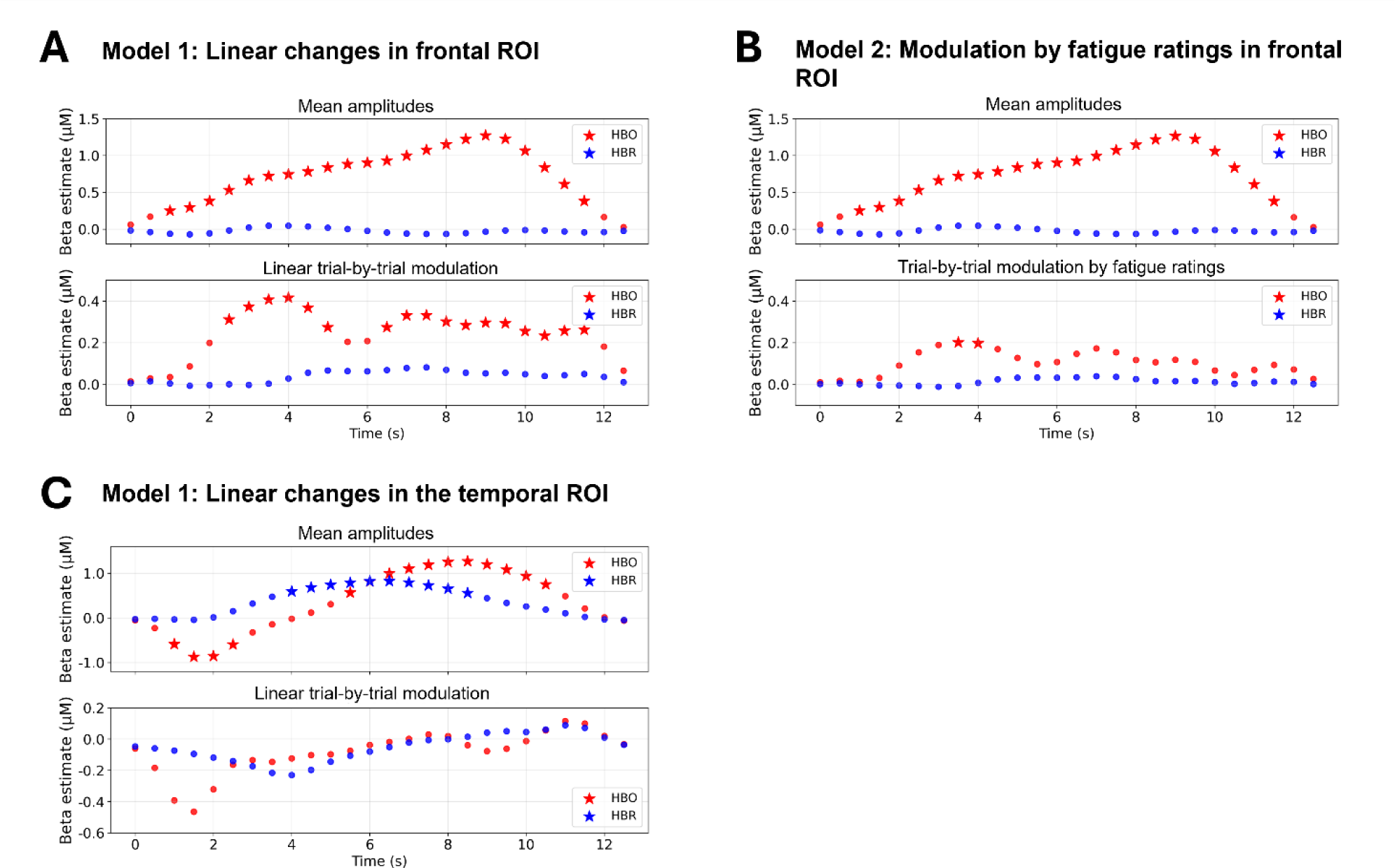
fNIRS analysis 2 – group level results. Group-level beta estimates for bins within the first 13 seconds of all accepted offer trials. Values were estimated using a linear mixed-effects models of the form: Beta ∼ Delay Bin (1-2c) × Chroma (HbO vs. HbR) × Condition (mean response vs. trial-by-trial modulation) + (1|participant) + (1|channel). **A.** Frontal ROI results from time-on-task model. **B.** Frontal ROI results from fatigue model. **C.** Temporal ROI results from time on task. In each panel, the upper plot shows mean response amplitudes for each delay bin in the respective ROI. The lower panel shows trial-by-trial modulations by either time on task or individual fatigue ratings, showing the strongest modulation of amplitudes over trials were linear modulations in the frontal ROI, at 4 seconds after offer onset. HbO is shown in red and HbR in blue. Significant beta values are indicated by stars.

As a control, the same model was repeated for the temporal ROI. The mean response was significant in several delay bins for both HbO and HbR (**Figure 12**C, upper panel). Both chromophores followed a similar response shape with an initial dip followed by a later peak. Importantly, no evidence for a linear modulation of response amplitude over the course of the experiment was observed in any delay bin (**Figure 12C**, lower panel).

In a final step, we tested whether not only time-on-task, but also individually reported fatigue moderated trial-by-trial amplitudes. Repeating the analysis with subjective fatigue ratings as the modulation regressor revealed significant beta values only for bin 8 and 9. Overall, the modulation time course showed a broadly similar shape to that observed for the time-on-task analysis (see **Figure 12B**, lower panel in comparison to **Figure 12A**, lower panel).

## Discussion

Listening-related fatigue in hearing aid users is believed to be one reason for withdrawal from conversations and social interactions, with subsequent consequences for hearing aid use, perceived hearing aid handicap, hearing aid satisfaction and overall well-being. Hypothesizing that fatigue accumulation does not depend on acoustic challenge alone, but also on how rewarding a situation is, we designed a lab-based listening game to study participants’ willingness to invest listening effort across different combinations of task difficulty and reward. Varying the difficulty levels (SNRs) and the number of reward points that could be earned for correctly repeating a sentence in noise, we investigated how hearing aid users decided whether or to accept or decline trial-by-trial offers to perform a listening task, how these choices varied as a function of time on task and self-reported fatigue, and whether these changes were related to frontal cortex hemodynamic activation. Overall, fatigue increased substantially over the course of the experiment, listening decisions reflected the expected effort–reward trade-off, and frontal activation during offer evaluation increased over time, particularly for demanding offers with relatively low rewards. However, time on task was more consistently associated with behavioral and neural changes than subjective fatigue ratings.

### Fatigue ratings reflected both task demands and listening success

Increasing fatigue ratings over the course of the experiment confirmed that prolonged engagement in effortful listening was associated with increasing subjective fatigue. Fatigue ratings increased on listening, but not on rest trials. Changes in ratings were also independently associated with listening difficulty and trial success, with larger increases on more difficult trials and on trials on which participants responded incorrectly. Moreover, both current-trial and previous-trial success contributed to changes in fatigue ratings, indicating that fatigue judgments were influenced not only by task demands but also by recent performance history and that these effects might be partially overlapping.

This pattern of results is consistent with previous reports of cognitive fatigue (Matthews et al., 2023), showing that fatigue ratings were sensitive not only to time on task and task demands, but also to whether participants successfully achieved the task goal. In line with Hockey (2013) and Davis et al. (2021), perceived failure to accurately repeat the sentence and thereby obtain the associated reward may reflect an unfavorable balance of invested effort and expected benefit, ultimately contributing to the experience of fatigue. However, it is also plausible that instead of necessarily *feeling* more fatigued, participants may have used their perceived success or the difficulty level of the trial as a heuristic for estimating their fatigue state. Frustration with one’s own performance may have also contributed to the reported fatigue ratings. Thus, the ratings may have reflected a broader negative affective response rather than fatigue exclusively.

Notably, the relationship between performance and fatigue ratings also varied with hearing status. Participants with less severe hearing loss showed a stronger association between trial-by-trial listening success and changes in fatigue ratings. One possible explanation is that listeners with greater hearing loss experience listening difficulties more frequently in everyday life, such that momentary successes or failures may carry less informational value when inferring their current level of fatigue.

In summary, we found strong evidence that listening success was associated with changes in fatigue ratings, while there was no evidence that offered reward or the number of reward points obtained on successful trials modulated Δ-fatigue. Likewise, although fatigue ratings increased with listening difficulty, exploratory analyses provided only limited evidence that actually exerted listening effort contributed more strongly than offered difficulty per se.

### Listen-versus-rest decisions

Despite reporting relatively high fatigue towards the end of the experiment, participants chose to rest on only 7% of trials on average, with some participants never selecting the rest option. This limited use of the rest option may have indicated that substantial fatigue was required before participants disengaged behaviorally. Alternatively, participants may have been reluctant to express fatigue behaviorally, possibly due to a desire to remain engaged or to be perceived as compliant with the task.

The decision model confirmed that participants behaved in accordance with the intended effort–reward structure of the paradigm. Participants were more likely to accept offers of easy and medium difficulty than hard offers. In addition, offers associated with a reward of 10 points were accepted more frequently than offers associated with 6 points, indicating that the point rewards provided effective motivation and influenced decision-making.

Furthermore, the probability of listening decreased with time on task, and subjective fatigue was also associated with listening decisions when considered separately. However, these effects did not interact with reward or difficulty, providing no evidence that the weighting of reward and difficulty changed as the experiment progressed or as participants reported greater fatigue. This differs from findings reported for cognitive fatigue by Matthew et al. (2023).

Finally, trial success also influenced subsequent choices. Following incorrect trials, participants were less likely to choose listening on the next trial. Together with the effects observed for fatigue ratings, these findings suggest that trial success contributed both to how participants evaluate their current fatigue state and to their willingness to accept subsequent listening offers.

### Frontal cortex activity associated with offer evaluation increases with time on task

A central aim of this study was to determine whether the accumulation of listening-related fatigue over the course of the experiment was accompanied not only by changes in subjective fatigue ratings and listen-versus-rest decisions, but also by changes in neural activity during offer evaluation. We found that frontal cortical activation during offer evaluation increased over the course of the experiment, particularly for high-difficulty offers associated with low and medium rewards.

These effects were confined to frontal regions and time windows linked to offer evaluation and were not observed in later phases of the trial or in temporal control regions. Together, these findings suggest that prolonged task engagement was associated with changes in neural processes involved in evaluating difficulty-reward trade-offs and determining whether a given offer was worth the anticipated effort.

Importantly, participants did not show stronger behavioral effort discounting over time: fatigue and time on task reduced the overall probability of accepting listening offers but did not alter how reward and difficulty were weighted in overt choices. In contrast, neural activity during offer evaluation was not constant across difficulty and reward conditions over time but increased more strongly for particularly demanding offers, especially those paired with lower rewards. One possibility is that participants maintained relatively stable decision policies despite increasing fatigue, but that doing so required greater frontal engagement as the experiment progressed. In this view, increased activation during offer evaluation may reflect additional or compensatory evaluative processing needed to sustain decision-making over time.

A subsequent analysis demonstrated that the increases in frontal cortical activity reflected a gradual linear increase in response amplitudes over time rather than only a categorical difference between the two experimental halves. Importantly, the strongest linear increases occurred in delay bins corresponding to the expected peak of the offer-related hemodynamic response shape, assuming a canonical response shape, supporting the definition of time windows in Analysis 1 and the conclusion that the observed effects were likely related to offer evaluation. Notably, while response amplitudes increased linearly with time on task, modulations by individual fatigue ratings showed a similar pattern across delay bins, albeit with smaller and largely non-significant beta values. Given the high correlation between fatigue ratings and time on task, both measures captured a similar underlying trend across the experiment. Nevertheless, time on task yielded more robust effects, suggesting that gradual changes associated with prolonged task engagement may provide a more sensitive account of the observed modulation in frontal offer-evaluation responses. Fatigue ratings, in contrast, were also influenced by trial difficulty and trial success, indicating that the additional state-dependent factors captured by fatigue ratings did not explain additional variability in frontal activity beyond that accounted for by time on task.

Given the established role of frontal brain regions in executive control and value-based decision making, increased frontal activation may reflect greater engagement of evaluative processes when deciding whether to accept particularly demanding listening offers. The generally high proportion of accepted offers suggests that participants initially tended to accept most offers. The increasing frontal cortical activation during evaluation of accepted offers with time on task may reflect growing hesitation about accepting them, that is, a more careful consideration of the option to rest. Such a shift may also have been accompanied by longer decision times, which could contribute to the larger hemodynamic responses observed later in the experiment. Future analyses of reaction times and gaze behavior during offer evaluation are needed to determine whether offer evaluation became more deliberate over time or remained similarly deliberate but became more cognitively demanding as the experiment progressed.

The precise functional interpretation of the frontal activation nonetheless remains challenging, given that the frontal ROI was large and the frontal cortex constitutes a functionally heterogeneous region involved in multiple high-level cognitive processes. Our ROI overlapped with regions associated with latent fatigue states derived from computational modelling in the fMRI work by Müller et al. (2021) on physical fatigue, suggesting that the observed activation may also reflect neural processes related to fatigue accumulation and the evaluation of the subjective value of listening effort. The increase in frontal activation may therefore reflect several partly overlapping mechanisms, including increased deliberation about whether an offer is worth accepting, compensatory recruitment to maintain stable decision-making despite fatigue, neural processing related to accumulated fatigue itself, or other executive control processes not specific to the effort-discounting paradigm.

Disentangling these mechanisms will require future studies using more spatially specific ROIs, individual-channel analyses, and computational models that separately estimate fatigue accumulation and subjective value. For example, distinct contributions of the left and right medial prefrontal cortex to decision-making processes have been suggested (Li et al., 2019). More spatially resolved analyses of subregions may also clarify the role of task difficulty, which did not show a systematic effect within the broad frontal ROI used. For example, cortical hemodynamic activity in left prefrontal regions has been linked to anticipation of task difficulty in both arithmetic (Vassena et al., 2019) and verbal-memory tasks (Song et al., 2026).

The effect of trial success on cortical activation may be a further factor worth investigating, given the observed evidence that performance shaped fatigue ratings and listening-versus-rest decisions. Advantageous versus disadvantageous decisions have been shown to elicit distinct frontal fNIRS responses in other decision-making paradigms, including the Iowa Gambling Task and moral decision-making tasks (Li et al., 2019; Vanutelli et al., 2020). In addition, during speech-in-noise listening, incorrect responses have been associated with higher PFC activation (Perron et al., 2025). It remains to be investigated whether successful versus unsuccessful sentence repetition may also influence subsequent effort valuation and the neural processing underlying fatigue-related listening decisions.

Finally, future studies may validate the present fNIRS findings using longer event durations and inter-stimulus intervals optimized for separating offer evaluation from later trial stages and by examining both HbO and HbR signals. In the present dataset, HbR showed limited variability and was therefore considered insufficiently sensitive to provide additional information beyond the HbO results.

Irrespective of whether the frontal signal primarily reflects fatigue accumulation directly, or decision processes that were moderated by fatigue, it is important that changes were observed during the evaluation of whether to engage in listening, before listening effort was actually exerted. Anticipatory decision periods may therefore provide useful information about fatigue-related changes in frontal cortical processing, complementing subjective fatigue ratings that can be noisy and context-dependent. Accordingly, future studies of fatigue build-up in hearing aid users should consider changes in frontal activation during decision processes, as these may be moderated by fatigue and could inform strategies to prevent fatigue-related symptoms in this population.

### Is time on task a better estimator of fatigue than subjective ratings?

Both listen-versus-rest decisions and trial-by-trial changes in frontal cortical responses during accepted offer evaluation were better predicted by trial number than by subjective fatigue ratings. However, this does not imply that time on task provides a more accurate measure of fatigue. Such an interpretation would require assuming that changes in choice behavior and frontal cortical activity represent ground-truth indicators of fatigue, whereas these measures may also reflect other processes associated with prolonged task engagement.

While subjective fatigue ratings may capture more nuanced moment-to-moment fluctuations in experienced fatigue, they are also subject to measurement limitations. Participants may have found it difficult to distinguish between nearby points on the 100-point scale, and ratings may additionally reflect a mixture of fatigue-related and other task-related states, including frustration, motivation, and engagement. Such factors may introduce variability that weakens associations with behavioral and neural measures. Moreover, the recruited hearing-aid users were experienced research participants and may have been accustomed to maintaining performance despite increasing fatigue, both in everyday life and in experimental settings, and therefore may have been less inclined to attend to or report developing listening-related fatigue.

Overall, we can therefore conclude that changes in listen-versus-rest decisions and frontal cortical activity unfolded gradually across the experiment and were more closely related to cumulative task exposure than to momentary self-reported fatigue.

While these remaining challenges in assessing momentary fatigue strengthen the motivation to develop indirect measures of fatigue, it should also be acknowledged that the usefulness of such measures may not depend on their ability to isolate fatigue as a single, specific construct. Measures that capture the gradual emergence of broader functional consequences of prolonged listening may still prove valuable for evaluating hearing-aid interventions and addressing complaints of listening-related fatigue.

## Conclusion

The present study investigated whether decisions to engage or disengage from listening, and associated frontal cortical activity are moderated by time on task or subjectively reported fatigue. Our findings suggest frontal hemodynamic responses during offer evaluation increased with time on task, particularly for demanding offers, even though behavioral reward-difficulty weighting remained relatively stable. Together, these findings suggest that processes associated with prolonged task engagement and accumulating fatigue are reflected during the evaluation of listening offers before listening effort is actually exerted.

In the long term, reliable markers associated with fatigue-build-up have the potential to support hearing rehabilitation by capturing aspects of sustained listening that are not currently assessed in routine clinical practice. Ultimately, such measures, together with a better understanding of the costs and benefits associated with listening, may support strategies that reduce exhaustion while promoting sustained engagement in meaningful and rewarding listening situations.

## Footnotes

1 The setup included a Tobii Pro Spectrum eye tracker and a Logitech C270 HD Webcam. An investigation of eye and facial features during sentence listening and their association with fatigue ratings and listen-versus-rest decisions is described in Dreneva et al. (in preparation; DOI to be added upon publication).

## Author contributions

A.S., C.C., D.W., I.J., M.A., T.D., and H.I. conceived the study. K.D., S.L., and M.A. contributed to the design of the listening effort-discounting paradigm. A.S. and C.C. led data acquisition. A.S. analyzed the data and wrote the manuscript. J.E. provided methodological input to the fNIRS analysis. A.S., D.W., and H.I. were primarily involved in data interpretation. All authors revised the manuscript and approved the final version.

## Acknowledgements

We would like to thank Alberte Hygum Valsted, Elisabeth Björk Schmidt, and Lena Havtorn for their support with participant recruitment, participant communication, and data collection, as well as for their feedback on the study design. We also thank Johannes Wienen, Anna Dreneva and Joseph Rovetti for valuable discussions regarding the study design, data collection, and analysis.

## Data availability

Summary data supporting the findings of this study along with scripts for group-level analysis are available in DTU Data and can be accessed during peer review via a private link. Upon publication, the data will be openly available in DTU Data at https://doi.org/10.11583/DTU.33393718.

## References

Alhanbali S, Dawes P, Lloyd S, et al. (2017) Self-Reported Listening-Related Effort and Fatigue in Hearing-Impaired Adults. Ear and hearing 38(1). Ear Hear: e39–e48.

Alhanbali S, Dawes P, Lloyd S, et al. (2018) Hearing Handicap and Speech Recognition Correlate With Self-Reported Listening Effort and Fatigue. Ear and hearing 39(3). Ear Hear: 470–474.

Alhanbali S, Dawes P, Millman RE, et al. (2019) Measures of Listening Effort Are Multidimensional. Ear and Hearing 40(5). Lippincott Williams and Wilkins: 1084–1097.

Bess FH and Hornsby BWY (2014) Commentary: listening can be exhausting--fatigue in children and adults with hearing loss. Ear and hearing 35(6). Ear Hear: 592–599.

Blümer M, Heeren J, Mirkovic B, et al. (2024) The Impact of Hearing Aids on Listening Effort and Listening-Related Fatigue - Investigations in a Virtual Realistic Listening Environment. Trends in Hearing 28. SAGE Publications Inc.

Brehm JW and Self EA (1989) The intensity of motivation. Annual review of psychology 40. Annu Rev Psychol: 109–131.

Burnham KP and Anderson DR (2004) Multimodel Inference: Understanding AIC and BIC in Model Selection Multimodel Inference Understanding AIC and BIC in Model Selection. 33. 261 Sociological Methods Research.

Cazzell M, Li L, Lin ZJ, et al. (2012) Comparison of neural correlates of risk decision making between genders: An exploratory fNIRS study of the Balloon Analogue Risk Task (BART). NeuroImage 62(3): 1896–1911.

Cumming J, Manchaiah V, Mahomed-Asmail F, et al. (2026) Understanding Hearing Aid Use and Nonuse of Adult Hearing Aid Recipients: A Qualitative Content Analysis. Journal of the American Academy of Audiology 37(1). American Academy of Audiology: 42.

Davis H, Schlundt D, Bonnet K, et al. (2021) Understanding Listening-Related Fatigue: Perspectives of Adults with Hearing Loss. International journal of audiology 60(6). Int J Audiol: 458–468.

Dimitrijevic A, Smith ML, Kadis DS, et al. (2019) Neural indices of listening effort in noisy environments. Scientific Reports 9(1). Nature Publishing Group: 11278.

Downs DW (1982) Effects of hearing and use on speech discrimination and listening effort. The Journal of speech and hearing disorders 47(2). J Speech Hear Disord: 189–193.

Glover GH (1999) Deconvolution of impulse response in event-related BOLD fMRI. NeuroImage 9(4). Academic Press Inc.: 416–429.

Gosselin PA and Gagné J-P (2010) Use of a dual-task paradigm to measure listening effort. Canadian Journal of Speech-Language Pathology and Audiology 34(1): 43–51.

Goutte C;, Nielsen FÅ and Hansen LK (2000) Modeling the hemodynamic response in fMRI using smooth FIR filters. I E E E Transactions on Medical Imaging 19(12). APA: 1188–1201.

Hamann A and Carstengerdes N (2023) Assessing the development of mental fatigue during simulated flights with concurrent EEG-fNIRS measurement. Scientific Reports 2023 13:1 13(1). Nature Publishing Group: 1–12.

Hockey R (2011) The psychology of fatigue: Work, effort and control. The Psychology of Fatigue: Work, Effort and Control. Cambridge University Press: 1–272.

Hockey R (2013) A motivation control theory of fatigue. The Psychology of Fatigue. Cambridge University Press: 132–154.

Holman JA, Drummond A and Naylor G (2020) The Effect of Hearing Loss and Hearing Device Fitting on Fatigue in Adults: A Systematic Review. Ear and Hearing 42(1). Wolters Kluwer Health: 1.

Holman JA, Drummond A and Naylor G (2021) Hearing Aids Reduce Daily-Life Fatigue and Increase Social Activity: A Longitudinal Study. Trends in Hearing 25. SAGE Publications Inc.

Hornsby BWY (2013) The effects of hearing aid use on listening effort and mental fatigue associated with sustained speech processing demands. Ear and hearing 34(5). Ear Hear: 523–534.

Hornsby BWY and Kipp AM (2016) Subjective Ratings of Fatigue and Vigor in Adults With Hearing Loss Are Driven by Perceived Hearing Difficulties Not Degree of Hearing Loss. Ear and hearing 37(1). Ear Hear: e1–e10.

Houben R, Van Doorn-Bierman M and Dreschler WA (2013) Using response time to speech as a measure for listening effort. International Journal of Audiology 52(11). Taylor & Francis: 753–761.

Ishii A, Tanaka M and Watanabe Y (2014) Neural mechanisms of mental fatigue. Reviews in the Neurosciences 25(4). Walter de Gruyter GmbH: 469–479.

Kahneman D and Tversky A (1973) On the psychology of prediction. Psychological Review 80(4): 237–251.

Keidser G, Dillon H, Mejia J, et al. (2013) An algorithm that administers adaptive speech-in-noise testing to a specified reliability at selectable points on the psychometric function. International Journal of Audiology 52(11). Taylor & Francis: 795–800.

Kressner AA (2023) Danish Sentence Test (DAST). Technical University of Denmark.

Kressner AA, Jensen-Rico KM, Kizach J, et al. (2024) A corpus of audio-visual recordings of linguistically balanced, Danish sentences for speech-in-noise experiments. Speech Communication 165. North-Holland: 103141.

Kressner AA, Jensen-Rico KM, Kofoed Pedersen A, et al. (2025) Psychoacoustic characterisation of linguistically balanced, Danish sentences for speech-in-noise experiments. International Journal of Audiology 64(11). Taylor and Francis Ltd.: 1129–1137.

Kressner AA, Jensen-Rico KM, Pedersen AK, et al. (2026) Validation of the Adaptive Danish Sentence Test (DAST): A Template-Based Approach for Developing Linguistically Rich Sentence-in-Noise Tests. Preprints. Epub ahead of print 24 April 2026. DOI: 10.20944/PREPRINTS202604.1764.V1.

Krueger M, Schulte M, Brand T, et al. (2017) Development of an adaptive scaling method for subjective listening effort. The Journal of the Acoustical Society of America 141(6). J Acoust Soc Am: 4680–4693.

Li Y, Chen R, Zhang S, et al. (2019) Hemispheric mPFC asymmetry in decision making under ambiguity and risk: An fNIRS study. Behavioural Brain Research 359. Elsevier: 657–663.

Lin CT, King JT, Chuang CH, et al. (2019) Exploring the Brain Responses to Driving Fatigue Through Simultaneous EEG and fNIRS Measurements. 10.1142/S0129065719500187 30(1). World Scientific Publishing Company.

Mackersie CL and Calderon-Moultrie N (2016) Autonomic nervous system reactivity during speech repetition tasks: Heart rate variability and skin conductance. Ear and Hearing 37. Lippincott Williams and Wilkins: 118S–125S.

Matthews J, Pisauro MA, Jurgelis M, et al. (2023) Computational mechanisms underlying the dynamics of physical and cognitive fatigue. Cognition 240. Elsevier B.V.

McGarrigle R, Munro KJ, Dawes P, et al. (2014) Listening effort and fatigue: What exactly are we measuring? A British Society of Audiology Cognition in Hearing Special Interest Group ‘white paper’. International Journal of Audiology 53(7). Informa Healthcare: 433–445.

McLaughlin DJ, Braver TS and Peelle JE (2021) Measuring the subjective cost of listening effort using a discounting task. Journal of Speech, Language, and Hearing Research 64(2). American Speech-Language-Hearing Association: 337–347.

Micula A, Flensborg-Madsen T and Christensen JH (2026) Influence of personality, cognition, and hearing loss on the association of real-world listening conditions with self-reported listening effort and heart rate. Hearing Research 475. Elsevier B.V.

Müller T and Apps MAJ (2019) Motivational fatigue: A neurocognitive framework for the impact of effortful exertion on subsequent motivation. Neuropsychologia 123. Elsevier Ltd: 141–151.

Müller T, Klein-Flügge MC, Manohar SG, et al. (2021) Neural and computational mechanisms of momentary fatigue and persistence in effort-based choice. Nature Communications 12(1). Nature Research.

Nihashi T, Ishigaki T, Satake H, et al. (2019) Monitoring of fatigue in radiologists during prolonged image interpretation using fNIRS. Japanese Journal of Radiology 37(6). Springer: 437–448.

Ohlenforst B, Zekveld AA, Lunner T, et al. (2017) Impact of stimulus-related factors and hearing impairment on listening effort as indicated by pupil dilation. Hearing Research 351. Elsevier: 68–79.

Peelle JE (2018) Listening effort: How the cognitive consequences of acoustic challenge are reflected in brain and behavior. Ear and Hearing 39(2): 204–214.

Perron M, Shatzer H, Zara M, et al. (2025) Age-related increased frontal activation in sentence comprehension reflects inefficiency, not compensation. Neurobiology of Aging 155. Elsevier: 100–112.

Pichora-Fuller MK, Kramer SE, Eckert MA, et al. (2016) Hearing Impairment and Cognitive Energy: The Framework for Understanding Effortful Listening (FUEL). Ear and hearing 37 Suppl 1. Ear Hear: 5S–27S.

Pollonini L, Bortfeld H and Oghalai JS (2016) PHOEBE: a method for real time mapping of optodes-scalp coupling in functional near-infrared spectroscopy. Biomedical Optics Express 7(12). The Optical Society: 5104.

Prokopiou PC, Xifra-Porxas A, Michalis Kassinopoulos ·, et al. (2022) Modeling the Hemodynamic Response Function Using EEG-fMRI Data During Eyes-Open Resting-State Conditions and Motor Task Execution. Brain Topography 35: 302–321.

Rakerd B, Seitz PF and Whearty M (1996) Assessing the cognitive demands of speech listening for people with hearing losses. Ear and hearing 17(2). Ear Hear: 97–106.

Richter M, Gendolla GHE and Wright RA (2016) Three Decades of Research on Motivational Intensity Theory: What We Have Learned About Effort and What We Still Don’t Know. Advances in Motivation Science 3. Elsevier: 149–186.

Rovetti J, Goy H, Nurgitz R, et al. (2021) Comparing verbal working memory load in auditory and visual modalities using functional near-infrared spectroscopy. Behavioural Brain Research 402. Elsevier B.V.

Rovetti J, Goy H, Zara M, et al. (2022) Reduced Semantic Context and Signal-to-Noise Ratio Increase Listening Effort As Measured Using Functional Near-Infrared Spectroscopy. Ear and Hearing 43(3). Lippincott Williams and Wilkins: 836–848.

Schneider EN, Bernarding C, Francis AL, et al. (2019) A Quantitative Model of Listening Related Fatigue. International IEEE/EMBS Conference on Neural Engineering, NER 2019-March. IEEE Computer Society: 619–622.

Shatzer HE and Russo FA (2023) Brightening the Study of Listening Effort with Functional Near-Infrared Spectroscopy: A Scoping Review. Seminars in Hearing 44(2). Thieme Medical Publishers, Inc.: 188–210.

Skau S, Bunketorp-Käll L, Kuhn HG, et al. (2019) Mental fatigue and functional near-infrared spectroscopy (fNIRS) – Based assessment of cognitive performance after mild traumatic brain injury. Frontiers in Human Neuroscience 13. Frontiers Media S.A.: 434384.

Skau S, Johansson B, Kuhn HG, et al. (2022) Segregation over time in functional networks in prefrontal cortex for individuals suffering from pathological fatigue after traumatic brain injury. Frontiers in Neuroscience 16. Frontiers Media S.A.

Song Y, Dahal P, Kim H ju, et al. (2026) Left frontal activity measured by fNIRS during a verbal memory task is modulated by predictability in cognitively healthy older adults. NeuroImage: Reports 6(3). Elsevier: 100380.

Vaisberg JM, Gilmore S, Qian J, et al. (2024) The Benefit of Hearing Aids as Measured by Listening Accuracy, Subjective Listening Effort, and Functional Near Infrared Spectroscopy. Trends in Hearing 28. SAGE Publications Inc.

Vanutelli ME, Meroni F, Fronda G, et al. (2020) Gender Differences and Unfairness Processing during Economic and Moral Decision-Making: A fNIRS Study. Brain Sciences 2020, Vol. 10, Page 647 10(9). Multidisciplinary Digital Publishing Institute: 647.

Varandas R, Lima R, Badia SBI, et al. (2022) Automatic Cognitive Fatigue Detection Using Wearable fNIRS and Machine Learning. Sensors 2022, Vol. 22, Page 4010 22(11). Multidisciplinary Digital Publishing Institute: 4010.

Vassena E, Gerrits R, Demanet J, et al. (2019) Anticipation of a mentally effortful task recruits Dorsolateral Prefrontal Cortex: An fNIRS validation study. Neuropsychologia 123. Pergamon: 106–115.

Wang Y, Kramer SE, Wendt D, et al. (2018) The Pupil Dilation Response During Speech Perception in Dark and Light: The Involvement of the Parasympathetic Nervous System in Listening Effort. Trends in Hearing 22. SAGE Publications Inc.

Yan Y, Guo Y and Zhou D (2025) Mental fatigue causes significant activation of the prefrontal cortex: A systematic review and meta-analysis of fNIRS studies. Psychophysiology. John Wiley and Sons Inc.

Yücel MA, Lühmann A v., Scholkmann F, et al. (2021) Best practices for fNIRS publications. Neurophotonics 8(01). International Society for Optics and Photonics: 012101.

Yücel MA, Luke R, Mesquita RC, et al. (2025) fNIRS reproducibility varies with data quality, analysis pipelines, and researcher experience. Communications Biology 2025 8:1 8(1). Nature Publishing Group: 1–17.

Zekveld AA, Kramer SE and Festen JM (2010) Pupil response as an indication of effortful listening: the influence of sentence intelligibility. Ear and hearing 31(4). Ear Hear: 480–490.

Zimeo Morais GA, Balardin JB and Sato JR (2018) FNIRS Optodes’ Location Decider (fOLD): A toolbox for probe arrangement guided by brain regions-of-interest. Scientific Reports 8(1). Nature Publishing Group: 1–11.

