## Supplementary Materials for "Fatigue-related changes in value-based decision-making during listening are associated with increased frontal cortical activation"

### Speech intelligibility performance

A linear mixed-effects model (LMM) indicated that performance decreased with increasing task difficulty ($F(2,145) = 234.83$, $p < .001$), whereas neither experimental half ($F(1,145) = 1.80$, $p = 0.18$) nor the Difficulty × Half interaction($F(2,145) = 1.58$, $p = .21$) significantly affected sentence repetition performance (**Figure S1**).

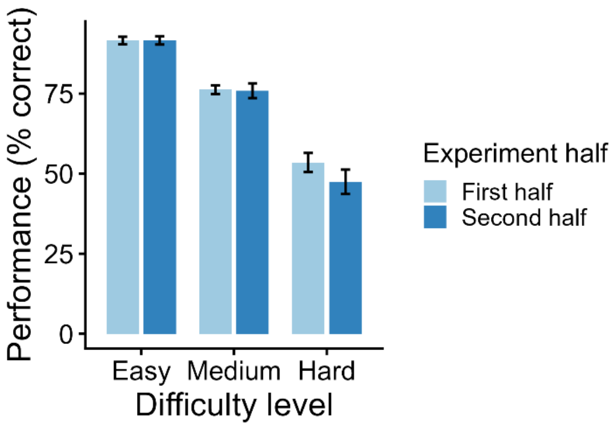


***Figure S1. Speech intelligibility performance.*** *Performance refers to % keywords repeated correctly and is shown for the three difficulty levels split into first and second experimental half respectively.*

### Supplementary analyses of non-retained models

#### Delta-fatigue model 1 (difficulty, reward, decision)

Delta-fatigue model 1 assessed the effect of various experimental variables on delta fatigue. The additive model was specified as delta_fatigue ~ difficulty + reward + decision + trial_num_z + PTA_z. The exploratory interaction model included all two-way interactions among these predictors: delta_fatigue ~ (trial_num_z + difficulty + reward + decision + PTA_z)^2. The improvement in fit was negligible once model complexity was taken into account ($\Delta AIC = 0.06$). The additive model was therefore retained for inference in the main manuscript. Additional results from the exploratory interaction model are reported here for completeness and transparency. Significant interaction terms in the interaction model included reward × decision and decision × PTA4 (Table S1, with the corresponding coefficient estimates reported in Table S3. Inspection of the model coefficients that the associations of reward and hearing loss with delta fatigue differed between listening and rest trials. Specifically, higher PTA4 values were associated with larger increases in delta fatigue during listening trials than during rest trials. PTA4 emerged as a significant predictor in the interaction model despite showing no overall association with delta fatigue in the additive model. This pattern may reflect the fact that the relationship between hearing loss and fatigue change was conditional on participants' listen-vs.-rest decisions. However, because the interaction model provided little improvement over the additive model, these findings should be interpreted as exploratory.

***Table S 1.*** *Type III ANOVA results for the exploratory two-way interaction model assessing effects of experimental variables on delta fatigue.*

| Effect | df | F | p |
|---|---|---|---|
| Trial number | 1 | 3.83 | .050 |
| Difficulty | 2 | 2.41 | .090 |
| Reward | 2 | 3.05 | .048* |
| Decision | 1 | 0.00 | .953 |
| PTA4 | 1 | 8.30 | .004** |
| Trial number × Difficulty | 2 | 2.49 | .083 |
| Trial number × Reward | 2 | 2.68 | .069 |
| Trial number × Decision | 1 | 0.90 | .342 |
| Trial number × PTA4 | 1 | 3.62 | .057 |
| Difficulty × Reward | 4 | 0.46 | .768 |
| Difficulty × Decision | 2 | 2.75 | .064 |
| Difficulty × PTA4 | 2 | 1.98 | .138 |
| Reward × Decision | 2 | 3.49 | .031* |
| Reward × PTA4 | 2 | 0.94 | .391 |
| Decision × PTA4 | 1 | 7.55 | .006* |

***Table S 2.*** *Coefficient estimates for the exploratory two-way interaction model assessing effects of experimental variables on delta fatigue.*

| Predictor | Estimate | SE | t | p |
|---|---|---|---|---|
| Intercept | 0.405 | 0.796 | 0.509 | .611 |
| Trial number | 0.624 | 0.319 | 1.958 | .050 |
| Difficulty: medium | −1.835 | 0.937 | −1.958 | .050 |
| Difficulty: hard | −0.562 | 0.811 | −0.693 | .488 |
| Reward: medium | 0.480 | 0.671 | 0.715 | .475 |
| Reward: high | −1.241 | 0.689 | −1.803 | .072 |
| Decision: listen | −0.047 | 0.795 | −0.059 | .953 |
| PTA4 | 0.741 | 0.257 | 2.880 | .004** |
| Trial number × Difficulty: medium | −0.169 | 0.161 | −1.055 | .292 |
| Trial number × Difficulty: hard | −0.366 | 0.164 | −2.231 | .026* |
| Trial number × Reward: medium | −0.046 | 0.161 | −0.288 | .773 |

| | | | | |
|---|---|---|---|---|
| Trial number × Reward: high | −0.345 | 0.161 | −2.150 | .032* |
| Trial number × Decision: listen | −0.276 | 0.290 | −0.950 | .342 |
| Trial number × PTA4 | −0.125 | 0.066 | −1.902 | .057 |
| Difficulty: medium × Reward: medium | 0.239 | 0.393 | 0.607 | .544 |
| Difficulty: hard × Reward: medium | 0.418 | 0.402 | 1.040 | .298 |
| Difficulty: medium × Reward: high | 0.394 | 0.392 | 1.003 | .316 |
| Difficulty: hard × Reward: high | 0.430 | 0.401 | 1.074 | .283 |
| Difficulty: medium × Decision: listen | 2.084 | 0.931 | 2.239 | .025* |
| Difficulty: hard × Decision: listen | 0.884 | 0.802 | 1.102 | .270 |
| Difficulty: medium × PTA4 | −0.228 | 0.161 | −1.418 | .156 |
| Difficulty: hard × PTA4 | −0.321 | 0.168 | −1.913 | .056 |
| Reward: medium × Decision: listen | −0.901 | 0.627 | −1.435 | .151 |
| Reward: high × Decision: listen | 0.889 | 0.648 | 1.372 | .170 |
| Reward: medium × PTA4 | −0.158 | 0.161 | −0.985 | .325 |
| Reward: high × PTA4 | 0.054 | 0.161 | 0.335 | .738 |
| Decision: listen × PTA4 | −0.603 | 0.219 | −2.747 | .006* |

***Table S 3. Delta fatigue model 1 – Estimated marginal means from the Difficulty × Decision interaction,*** *included to explore whether fatigue was more closely related to offered difficulty or effort exertion. Although the overall interaction did not reach significance, the estimated means suggested larger listen-rest differences for medium and hard than for easy difficulty offers.*

| Offered difficulty | Rest mean (95% CI) | Listen mean (95% CI) | Δ (Listen – Rest) | p |
|---|---|---|---|---|
| Easy | 0.15 (-1.26, 1.56) | 0.10 (-0.12, 0.33) | -0.05 | .944 |
| Medium | -1.47 (-2.61, -0.33) | 0.56 (0.33, 0.79) | +2.03 | < .001** |
| Hard | -0.13 (-0.76, 0.50) | 0.71 (0.46, 0.95) | +0.83 | .016* |

### Delta fatigue model 2 (response accuracy)

Delta fatigue model 2 assessed the effect of response accuracy on the trials change in fatigue rating. The additive model was specified as *delta_fatigue ~ trial_num_z + difficulty + reward + trial success + PTA_z*. The exploratory interaction model included all two-way interactions among these predictors: *delta_fatigue ~ (trial_num_z + difficulty + reward + trial success + PTA_z)^2*. The interaction model showed substantially poorer fit than the additive model once model complexity was taken into account (ΔAIC = 5.08). The additive model was therefore retained for inference in the main manuscript, while additional results from the exploratory interaction model are reported here for completeness and transparency. The exploratory interaction model revealed a significant trial success × PTA4 interaction (Table S3; coefficients in Table S4). Inspection of the coefficient estimates suggested that increasing PTA4 was associated with smaller differences in delta fatigue between correct and incorrect listening trials, but larger differences between correct listening trials and rest trials.

***Table S4.*** *Type III ANOVA results for the exploratory two-way interaction model assessing effects of trial success on delta fatigue.*

| Effect | df | F | p |
|---|---|---|---|
| Trial number | 1 | 5.81 | .016* |
| Difficulty | 2 | 0.31 | .732 |
| Reward | 2 | 1.17 | .310 |
| Correctness | 2 | 2.42 | .089 |
| PTA4 | 1 | 0.95 | .331 |
| Trial number × Difficulty | 2 | 2.20 | .111 |
| Trial number × Reward | 2 | 2.64 | .072 |
| Trial number × Correctness | 2 | 0.44 | .641 |
| Trial number × PTA4 | 1 | 3.78 | .052 |
| Difficulty × Reward | 4 | 0.65 | .627 |
| Difficulty × Correctness | 4 | 1.75 | .136 |
| Difficulty × PTA4 | 2 | 0.66 | .515 |
| Reward × Correctness | 4 | 2.09 | .079 |
| Reward × PTA4 | 2 | 0.94 | .392 |
| Correctness × PTA4 | 2 | 5.92 | .003** |

***Table S5.*** *Coefficient estimates for the exploratory two-way interaction model assessing effects of trial success on delta fatigue.*

| Predictor | Estimate | SE | t | p |
|---|---|---|---|---|
| Intercept | 0.310 | 0.198 | 1.566 | .117 |
| Trial number | 0.349 | 0.145 | 2.410 | .016* |
| Difficulty: medium | 0.204 | 0.285 | 0.715 | .475 |
| Difficulty: hard | 0.010 | 0.319 | 0.032 | .974 |
| Reward: medium | -0.391 | 0.278 | -1.408 | .159 |
| Reward: high | -0.339 | 0.277 | -1.222 | .222 |
| Correctness: incorrect | 1.698 | 0.772 | 2.201 | .028* |
| Correctness: not tested | 0.096 | 0.798 | 0.120 | .904 |
| PTA4 | 0.143 | 0.147 | 0.973 | .331 |
| Trial number × Difficulty: medium | -0.166 | 0.162 | -1.026 | .305 |
| Trial number × Difficulty: hard | -0.367 | 0.175 | -2.097 | .036* |
| Trial number × Reward: medium | -0.048 | 0.161 | -0.297 | .767 |
| Trial number × Reward: high | -0.343 | 0.160 | -2.135 | .033* |
| Trial number × Correctness: incorrect | -0.018 | 0.195 | -0.093 | .926 |
| Trial number × Correctness: not tested | 0.268 | 0.295 | 0.906 | .365 |
| Trial number × PTA4 | -0.128 | 0.066 | -1.944 | .052 |
| Difficulty: medium × Reward: medium | 0.262 | 0.397 | 0.661 | .509 |
| Difficulty: hard × Reward: medium | 0.575 | 0.428 | 1.341 | .180 |
| Difficulty: medium × Reward: high | 0.424 | 0.394 | 1.076 | .282 |
| Difficulty: hard × Reward: high | 0.525 | 0.427 | 1.229 | .219 |
| Difficulty: medium × Correctness: incorrect | -0.918 | 0.803 | -1.143 | .253 |
| Difficulty: hard × Correctness: incorrect | -0.755 | 0.773 | -0.977 | .329 |
| Difficulty: medium × Correctness: not tested | -2.002 | 0.931 | -2.151 | .032* |
| Difficulty: hard × Correctness: not tested | -0.603 | 0.807 | -0.748 | .455 |
| Difficulty: medium × PTA4 | -0.175 | 0.163 | -1.074 | .283 |

| | | | | |
|---|---|---|---|---|
| Difficulty: hard × PTA4 | -0.160 | 0.178 | -0.901 | .368 |
| Reward: medium × Correctness: incorrect | -0.510 | 0.482 | -1.058 | .290 |
| Reward: high × Correctness: incorrect | -0.204 | 0.492 | -0.415 | .678 |
| Reward: medium × Correctness: not tested | 0.775 | 0.642 | 1.209 | .227 |
| Reward: high × Correctness: not tested | -0.971 | 0.659 | -1.474 | .141 |
| Reward: medium × PTA4 | -0.163 | 0.161 | -1.014 | .311 |
| Reward: high × PTA4 | 0.046 | 0.161 | 0.288 | .774 |
| Correctness: incorrect × PTA4 | -0.443 | 0.208 | -2.133 | .033* |
| Correctness: not tested × PTA4 | 0.480 | 0.226 | 2.127 | .034* |
| Reward: high × Correctness: incorrect | -0.204 | 0.492 | -0.415 | .678 |
| Reward: medium × Correctness: not tested | 0.775 | 0.642 | 1.209 | .227 |
| Reward: high × Correctness: not tested | -0.971 | 0.659 | -1.474 | .141 |
| Reward: medium × PTA4 | -0.163 | 0.161 | -1.014 | .311 |
| Reward: high × PTA4 | 0.046 | 0.161 | 0.288 | .774 |
| Correctness: incorrect × PTA4 | -0.443 | 0.208 | -2.133 | .033* |
| Correctness: not tested × PTA4 | 0.480 | 0.226 | 2.127 | .034* |

### Delta fatigue model 3 (response accuracy versus feedback)

Delta fatigue model 3 assessed whether trial-by-trial changes in fatigue ratings were more strongly associated with current-trial response accuracy or previous-trial response accuracy (i.e., feedback from the preceding trial). The additive model was specified as *delta_fatigue ~ difficulty + reward + trial success + lag_difficulty + lag_reward + lag_trial success + PTA_z*. The exploratory interaction model additionally included theoretically motivated interactions involving current- and previous-trial response accuracy: *trial success × PTA_z*, *lag_trial success × PTA_z*, *trial success × difficulty*, and *lag_trial success × difficulty*. The interaction model provided a better fit to the data than the additive model once model complexity was taken into account (ΔAIC = 4.80) and was therefore retained for inference in the main manuscript. Results from the additive model are reported here for completeness and transparency. In the additive model, current-trial response accuracy predicted delta fatigue, with incorrect trials associated with a 0.71-unit larger increase in fatigue ratings than correct trials, whereas previous-trial response accuracy, reward, and PTA4 did not explain additional variance (Table S5, coefficients in Table S6).

***Table S6.*** *Type III ANOVA results the additive model testing effects current versus previous trial's response accuracy on delta fatigue. The model was restricted to listening trials that were preceded by another listening trial.*

| **Effect** | **df** | **F** | **p** |
|---|---|---|---|
| Difficulty | 2 | 2.98 | .051 |
| Reward | 2 | 1.03 | .358 |
| Current-trial response accuracy | 1 | 12.68 | < .001** |
| Previous-trial response accuracy | 1 | 0.16 | .691 |
| Previous-trial difficulty | 2 | 1.13 | .322 |
| Previous-trial reward | 2 | 0.12 | .890 |
| PTA4 | 1 | 0.06 | .807 |

***Table S7***. *Coefficient estimates for the additive model testing effects current versus previous trial's response accuracy on delta fatigue.*

| **Predictor** | **Estimate** | **SE** | **t** | **p** |
|---|---|---|---|---|
| Intercept | 0.354 | 0.214 | 1.652 | .099 |
| Difficulty: medium | 0.381 | 0.168 | 2.269 | .023 |
| Difficulty: hard | 0.346 | 0.182 | 1.906 | .057 |
| Reward: medium | -0.232 | 0.167 | -1.389 | .165 |
| Reward: high | -0.065 | 0.166 | -0.391 | .696 |
| Accuracy: incorrect | 0.710 | 0.199 | 3.560 | < .001** |
| Previous difficulty: medium | -0.168 | 0.168 | -0.999 | .318 |
| Previous difficulty: hard | -0.266 | 0.181 | -1.474 | .141 |
| Previous reward: medium | 0.006 | 0.167 | 0.038 | .969 |
| Previous reward: high | -0.066 | 0.166 | -0.395 | .693 |
| Previous accuracy: incorrect | -0.080 | 0.200 | -0.398 | .691 |
| PTA4 | -0.017 | 0.072 | -0.244 | .807 |

### Decision models

***Table S8***. *Type III Wald $\chi^2$ tests for the follow-up decision model examining whether time on task altered how participants weighed task difficulty and reward when deciding whether to listen or rest: decision ~ difficulty * trial_num_z * reward + (1 | subject_idx)*

| **Effect** | **$\chi^2$** | **df** | **p** |
|---|---|---|---|
| (Intercept) | 91.22 | 1 | < .001 *** |
| Difficulty | 63.60 | 2 | < .001 *** |
| Trial number | 0.47 | 1 | .494 |
| Reward | 0.89 | 2 | .641 |
| Difficulty × Trial Number | 0.52 | 2 | .771 |
| Difficulty × Reward | 0.42 | 4 | .981 |
| Trial Number × Reward | 1.34 | 2 | .512 |
| Difficulty × Trial Number × Reward | 2.32 | 4 | .678 |

***Table S9.*** *Type III Wald $\chi^2$ tests for the follow-up decision model examining whether subjective fatigue altered how participants weighed task difficulty and reward when deciding whether to listen or rest: decision ~ difficulty * fatigue_z * reward + (1 | subject_idx).*

| **Effect** | **$\chi^2$** | **df** | **p** |
|---|---|---|---|
| Difficulty | 64.80 | 2 | < .001*** |
| Fatigue | 0.02 | 1 | .886 |
| Reward | 1.11 | 2 | .574 |
| Difficulty × Fatigue | 1.00 | 2 | .607 |
| Difficulty × Reward | 0.34 | 4 | .987 |
| Fatigue × Reward | 1.66 | 2 | .435 |
| Difficulty × Fatigue × Reward | 3.57 | 4 | .467 |

### fNIRS models

***Table S10***. *Type III ANOVA results for the frontal ROI- early-window model predicting frontal HbO concentration during the early offer-evaluation time window. Fixed effects were tested for the model: HbO ~ Reward × Difficulty × Half + fatigability + (1 | Participant) + (1 | Channel). Reward (6, 8, 10 points), Difficulty (easy, medium, hard), and Half (first, second) were treatment-coded. Fatigability was quantified as the participant-specific slope of fatigue ratings over time and z-scored across participants.*

| **Effect** | **df** | **F** | **p** |
|---|---|---|---|
| Reward | 2 | 0.13 | .870 |
| Difficulty | 2 | 5.15 | .006** |
| Half | 1 | 55.9 | <.001*** |
| Fatigability | 1 | 0.18 | .67 |
| Reward × Difficulty | 4 | 10.00 | <.001*** |
| Reward × Half | 2 | 3.78 | .023* |
| Difficulty × Half | 2 | 34.91 | <.001*** |
| Reward × Difficulty × Half | 4 | 8.73 | <.001*** |

***Table S11***. *Type III ANOVA results for the follow-up model: HbO ~ ROI * window* ROI* Half + (1 | Participant) + (1 | Channel).*

| **Effect** | **df** | **F** | **p** |
|---|---|---|---|
| Window | 1 | 110.96 | < .001*** |
| ROI | 1 | 44.69 | < .001*** |
| Half | 1 | 5.69 | .017* |
| Window × ROI | 1 | 23.92 | < .001*** |
| Window × Half | 1 | 1.33 | .248 |
| ROI × Half | 1 | 4.44 | .035* |
| Window × ROI × Half | 1 | 3.79 | .052 |

***Table S12***. *Estimated marginal means of HbO concentration (μM) for each combination of ROI, time window, and experimental half, derived from the ROI × Window × Half mixed-effects model. The final column shows pairwise contrasts comparing the first and second half of the experiment within each ROI and time window. Significant half-related increases in activation were observed only in the early frontal time window. P-values for the pairwise contrasts are Bonferroni-corrected.*

| **ROI** | **Time window** | **Half 1 Mean (95% CI)** | **Half 2 Mean (95% CI)** | **Δ (Half 2 – Half 1)** | **p** |
|---|---|---|---|---|---|
| Frontal | Early | 0.41 (-0.47, 1.29) | 0.99 (0.10, 1.87) | +0.57 | < .001** |
| Temporal | Early | -0.82 (-1.79, 0.15) | -0.86 (-1.83, 0.12) | -0.04 | .83 |
| Frontal | Late | 1.81 (0.93, 2.69) | 1.92 (1.04, 2.80) | +0.11 | .35 |

| | | | | | |
|---|---|---|---|---|---|
| Temporal | Late | -0.45 (-1.43, 0.52) | -0.37 (-1.35, 0.60) | -0.08 | .66 |

***Table S13.*** *Type III ANOVA results for the short-channel control model. The model was specified as HbO ~ reward × difficulty × half + (1 | participant) and tested whether corresponding half-related effects were present in extracerebral hemodynamic signals. No main effect of half or interactions involving half were observed.*

| Effect | df | F | p |
|---|---|---|---|
| Reward | 2 | 3.43 | .033* |
| Difficulty | 2 | 0.48 | .620 |
| Half | 1 | 0.33 | .568 |
| Reward × Difficulty | 4 | 0.64 | .634 |
| Reward × Half | 2 | 0.05 | .952 |
| Difficulty × Half | 2 | 0.51 | .600 |
| Reward × Difficulty × Half | 4 | 1.28 | .276 |
